# Anatomy, transcription dynamics and evolution of wheat ribosomal RNA loci deciphered by a multi-omics approach

**DOI:** 10.1101/2020.08.29.273623

**Authors:** Zuzana Tulpová, Aleš Kovařík, Helena Toegelová, Pavla Navrátilová, Veronika Kapustová, Eva Hřibová, Jan Vrána, Jiří Macas, Jaroslav Doležel, Hana Šimková

**Author notes:** **Correspondence**: Hana Šimková.

## Abstract

**Background and Aims:** Three out of four RNA components of ribosomes are encoded by 45S rDNA loci, whose transcripts are processed into 18S, 5.8S and 26S ribosomal RNAs. The loci are organized as long head-to-tail tandem arrays of nearly identical units spanning over several megabases of sequence. Due to this peculiar structure, the number of rRNA genes, their sequence composition and expression status remain unclear, especially in complex polyploid genomes harbouring multiple loci. Here we conducted a complex study to decipher structure and activity of both major and minor rRNA loci in hexaploid bread wheat (*Triticum aestivum*).

**Methods:** We employed an original, multi-omics approach, combining chromosome flow sorting and optical mapping with transcriptome and methylome sequencing.

**Key Results:** The former two techniques enabled unbiased quantification of rDNA units in particular loci of the wheat genome. Total number of rRNA genes organized in tandem arrays was 4388, with 64.1, 31.4, 3.9 and 0.7% located in short arms of chromosomes 6B, 1B, 5D and 1A, respectively. At the expression level, only 1B and 6B loci contributed to transcription at roughly 2:1 ratio. The 1B:6B ratio varied among five analysed tissues (embryo, coleoptile, root tip, primary leaf, mature leaf), being the highest (2.64:1) in mature leaf and lowest (1.72:1) in coleoptile. Cytosine methylation was considerably higher in CHG contexts in the silenced 5D locus compared to the active 1B and 6B loci.

**Conclusions:** A fine genomic organization and tissue-specific expression of rRNA loci were deciphered, for the first time, in a complex polyploid species. We documented various mechanisms of rRNA dosage control, including gene elimination and stable inactivation related to nucleolar subdominance of A and D-genome loci, and a subtle, developmentally regulated silencing of one of the major loci. The results are discussed in the context of wheat evolution and transcription regulation.

## INTRODUCTION

Ribosomal DNA (rDNA) is a general term for genes encoding several types of ribosomal RNA (rRNA), which are crucial components of ribosomes. 45S rDNA locus, specific for eukaryotic genomes, is associated with the nucleolus and is termed nucleolus organizer region (NOR). 5S rRNA gene loci are located apart from the NOR locus in a majority of organisms. Both nuclear 5S and 45S rDNA units are mostly organized as long head-to-tail tandem arrays. The 45S rDNA unit is composed of conserved genes for 18S, 5.8S and 26S rRNA, separated by internal transcribed spacers ITS1 and ITS2. A smaller part of the unit is a non-transcribed intergenic spacer (IGS) composed of several types of repeats of variable size and sequence. The 45S rDNA is transcribed by polymerase I into a large precursor, which is processed into the three types of rRNA. The 5S rRNA genes are approximately 120 bp long and, interspersed by non-transcribed spacers of a few hundred base pairs, they form arrays spanning over several kilobases, which makes them accessible to long-read sequencing technologies such as PacBio (Symonová *et al*., 2017). On the contrary, 45S rDNA unit size ranges from 8 to13 kb across yeast, plant and animal genomes (Gerlach and Bedbrook, 1979; Pruitt and Meyerowitz, 1986; Nelson *et al*., 2019) and its arrays span megabases of sequence. This precludes their complete assembling from whole-genome next-generation-sequencing data and was only possible when coupling clone-by-clone approach with long-read nanopore sequencing (Sims *et al*., 2021).

The absence of the 45S rDNA arrays from genome assemblies hampers their analysis, which is especially challenging in complex and polyploid genomes comprising multiple rRNA gene loci, as is the case of bread wheat. Bread wheat (*Triticum aestivum* L.) is an allohexaploid species with a genome of 16.58 Gb (Doležel *et al*., 2018), composed of three subgenomes (2n=6x=42, AABBDD). The huge genome size, polyploidy and, consequently, a lack of a genome sequence hampered genome analysis for a long time. On the other hand, the large chromosomes and complex genome composition attracted the attention of cytogeneticists and stimulated research on wheat NORs by microscopic and other techniques. These studies made use of a collection of aneuploid lines generated for *T. aestivum* cv. Chinese Spring (Sears 1954) that enabled assigning NORs to particular chromosomes and studying their transcription by cytogenetic techniques. As early as in late 1950’s, Crosby (1957) reported that there were at least four different chromosomes in hexaploid wheat, 1A, 1B, 6B and 5D, able to form nucleoli. In *T. aestivum* ‘Chinese Spring’ (CS), large nucleoli were assigned to chromosomes 1B and 6B, while smaller ones, micronucleoli, were associated with other chromosomes, mostly the 5D (Darvey and Driscoll, 1972). Nevertheless, the activity of the minor loci has not been broadly confirmed.

First estimates of numbers of rRNA gene copies contained within the bread wheat genome were done by Flavell and co-workers (summarized in Flavell and O’Dell, 1976). Using rRNA/DNA hybridization assays, they estimated 9150 copies of 18S-25S rRNA genes per 2C nucleus of CS wheat, out of which 5500 (60%), 2700 (30%) and 950 (10%) were assigned to 6B, 1B and other chromosomes (predominantly 5D and 1A), respectively. Surprisingly, rRNA gene number in 1B and 6B correlated negatively with nucleolar volume in root tip cells of CS wheat where the volume of nucleolus on chromosome 1B was twice of that on chromosome 6B (Martini and Flavell, 1985), indicating double transcription activity of the 1B NOR compared to that of 6B. This study also demonstrated that the minor NORs on 5D and 1A increased their activity when the major NORs were deleted. In the most recent study on wheat NORs, Handa *et al*. (2018) used quantitative PCR (qPCR) and fluorescence *in situ* hybridization to quantify rDNA units in the bread wheat genome. Their estimate for the total copy number of rDNA units per CS wheat genome (1C) was 11,160 (100 Mb sequence), which was more than double amount compared to the study by Flavell and O’Dell, 1976. Besides, the authors compared rDNA units extracted from the reference genome of bread wheat, the International Wheat Genome Sequencing Consortium (IWGSC) RefSeq v1.0 (IWGSC, 2018), and identified four rDNA subtypes and analysed their expression, which was not proportional to their representation in the genome. They also positioned the major 45S rDNA loci on chromosomes 1B and 6B, taking the presence of rDNA clusters in the sequence as a clue.

Our BLAST search for 45S rDNA in the IWGSC RefSeq v1.0 genome, carried out in Ensembl Plants (https://plants.ensembl.org/Triticum_aestivum/Tools/Blast), revealed several additional clusters of rDNA (Fig. 1A), mainly on short arms of chromosomes 1A (1AS), 1B (1BS), 5D (5DS), 7D (7DS) and long arms of 1B (1BL) and 4A (4AL), most of which likely correspond to minor loci identified previously by *in situ* hybridization (Mukai *et al*., 1991). However, none of the clusters in the IWGSC RefSeq, including those assigned to *Nor-B1* and *Nor-B2*, comprised a regular array of rDNA units. This virtual absence of genomic information on the bread wheat NORs, together with discrepancies in rDNA copy-number estimates and only vague information on transcriptional activity of particular loci, limited mostly to root tissues investigated by the cytogeneticists, call for a detailed study to resolve sequence organization and tissue-specific expression of rRNA genes from individual loci.

**Fig 1.**
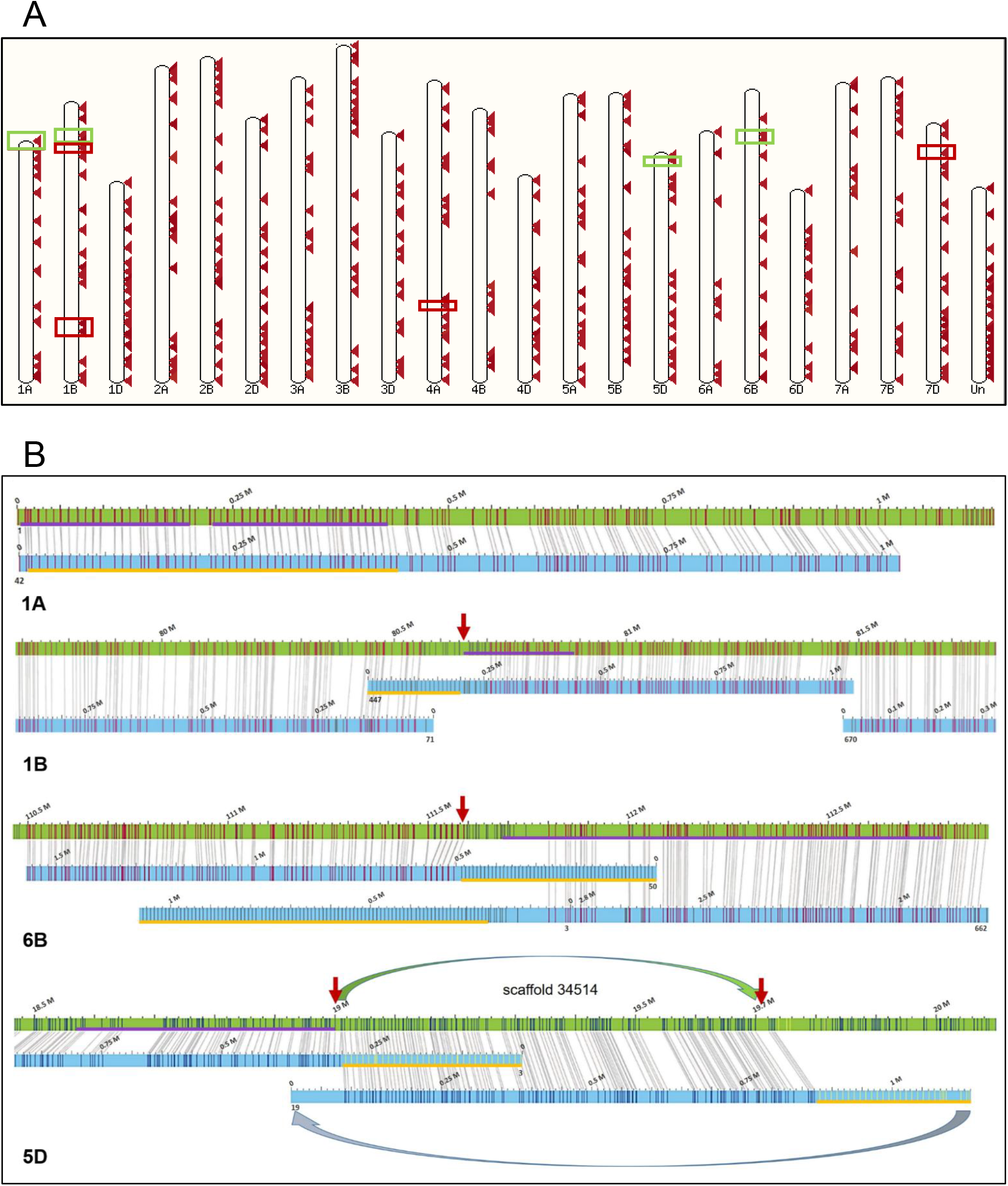
45S rDNA loci in bread wheat genome. (A) Graphical outcome of a BLAST search in Ensembl Plants/Triticum aestivum database for sequences homologous to wheat consensus rDNA unit in the IWGSC RefSeq v1.0 genome. The search was done with default settings. Vast majority of hits correspond to rDNA fragments; positions of larger rDNA clusters are marked by rectangles, with green colour pointing to rDNA loci analysed in the study. About 30% hits fall into unassigned scaffolds (Un). (B) Positioning of 45S rDNA arrays by optical mapping. Optical maps (blue bars) generated from rDNA-bearing chromosomes 1AS, 1BS, 6BS and 5DS were aligned to corresponding pseudomolecules of IWGSC RefSeq v1.0 wheat genome (green bars), digested *in silico* with enzymes used to construct the optical map. The vertical lines represent BspQI (red) and DLE-1 (blue) recognition sites labelled in particular optical map. Regular pattern with 8.4-9.6-kb spacing (highlighted by a yellow line) indicates presence of a 45S rDNA array. Only the irregular rDNA array in 1AS is present in the sequence while the regular arrays in the other chromosomes are missing in the pseudomolecules. The relevance of the arrays for the rDNA is supported by the presence of rDNA fragments found in the adjacent genomic regions (highlighted by violet lines). Sequence scaffold 34514, flanking the 5DS rDNA array at the proximal side, is misoriented in the IWGSC RefSeq v1.0 genome. The red arrows indicate scaffold boundaries in the reference genome.

We showed previously (Kapustová *et al*., 2019) that Bionano optical mapping, a technology that labels and visualizes short sequence motifs along 150-kb to 1-Mb long DNA molecules (Lam *et al*., 2012), is suitable for identification of rDNA arrays in a genomic context thanks to the presence of the labelled motifs in the 45S rDNA unit. Here, we used this technology, combined with separation of particular rDNA-bearing wheat chromosome arms by flow cytometry (Staňková *et al*., 2016), to search for the presence of rDNA arrays in the anticipated positions in 1BS and 6BS chromosome arms as well as in 1AS and 5DS, which are expected to bear minor rDNA loci. Additionally, analysis of optical-map raw data for the occurrence of tandemly organized repeats provided a novel approach to quantify rDNA units that discriminates potentially functional units organized in arrays from dispersed and fragmented ones, which are likely non-functional. Engagement of the major and the minor loci in rRNA synthesis was assessed in five tissues by analysing RNA sequencing data for chromosome-specific sequence variants in 26S rRNA and the transcriptional activity of particular loci was compared with cytosine methylation level in the rDNA units. Moreover, the high-throughput RNA-seq data provided information on the dynamic contribution of particular organelles to the overall rRNA production. Based on the data obtained, we outline a scenario for the evolutionary histories of rDNA loci in three wheat subgenomes and propose a model for control of rRNA amount through genetic and epigenetic mechanisms.

## MATERIALS AND METHODS

### Plant material

Seeds of double ditelosomic stocks of bread wheat (*Triticum aestivum*, L.) cv. Chinese Spring, dDt1AS, dDt1BS, dDt6BS and dDt5DS (Sears and Sears, 1978), were kindly provided by Prof. Bikram Gill (Kansas State University, Manhattan, USA). Seeds of *T. aestivum* cv. Chinese Spring with a standard karyotype were provided by Dr. Pierre Sourdille (INRAE, Clermont-Ferrand, France).

### Bionano optical mapping

Chromosome arms 1AS, 1BS, 6BS and 5DS were purified in million amounts by flow sorting from corresponding telosomic lines according to Kubaláková *et al*. (2002) and used for HMW DNA preparation as described in Staňková *et al*. (2016). Bionano optical maps (OMs) of 1AS, 1BS and 6BS were constructed using NLRS labelling chemistry (Nt.*BspQI* enzyme) and data were generated on the Irys platform (Bionano Genomics, San Diego, USA). The 5DS arm was labelled by DLS chemistry (DLE-1 enzyme) and molecules were analysed on the Saphyr platform (Bionano Genomics). DNA labelling and data collection followed protocols of Bionano Genomics (https://bionanogenomics.com/products/bionano-prep-kits/) with minor modifications. Optical map of the 1AS was assembled by Bionano IrysSolve 2.1.1 software, using optArguments_human.xml file. Assemblies of 1BS, 6BS and 5DS OMs were done by Bionano Solve 3.4 applying cmaps generated from fasta files of 1BS, 6BS and 5DS pseudomolecules of the IWGSC RefSeq v1.0 (IWGSC, 2018) as a reference. Assembly parameters specified by optArguments_nonhaplotype_noEs_noCut_irys.xml file were used for 1BS and 6BS map assembly whereas optArguments_nonhaplotype_noEs_noCut_DLE_saphyr.xml file were used for the 5DS OM. Information about input DNA amount, labelling chemistry, quantity and quality of data used for the OM assembly are stated in Table S1 **[Supplementary Information]**. The optical maps were aligned to the wheat reference using Bionano Solve v3.4 with RefAligner v9232 and pValue threshold of 1e−15 was applied for the query-to-anchor comparison. The alignments were visualized in Bionano Access v1.5. The OMs of 1BS and 6BS showed good concordance between distances in the map and in the sequence (length differences <1%), while alignment of the 5DS map revealed ∼8% expansion of the map compared to the reference sequence (Fig. 1B). To eliminate this inaccuracy, coefficient of 0.92 was used to correct the rDNA unit size estimate from the OM data.

### Quantification of units in rDNA arrays

An algorithm included in the IrysView 2.0 software package (Bionano Genomics) was used to identify labelled tandemly organized repeats in size-filtered (>150 kb) raw (single-molecule) data. Regular arrays of six and more repeat units were considered in the analysis. Repeat stretch tolerance of 0.1 was applied for 5DS (one DLE1 site per unit) while more relaxed 0.19 tolerance was used for 1BS and 6BS data generated with Nt.BspQI enzyme, because of two merging recognition sites in the unit. The detected repeats were quantified and their unit size and frequency in the dataset were plotted in a histogram for visual analysis. The fractions of telocentric chromosomes purified by flow cytometric sorting were contaminated by a mixture of other chromosomes/arms, totalling 9.6-19.7% of the sorted particles (Table 1). This was considered in recalculating the proportion of rDNA units per pure chromosomal fraction. Since the 1BS, 6BS and 5DS arms differ significantly in size (Table 1), their mutual contamination was unlikely and was not considered in the recalculation.

**Table 1.**
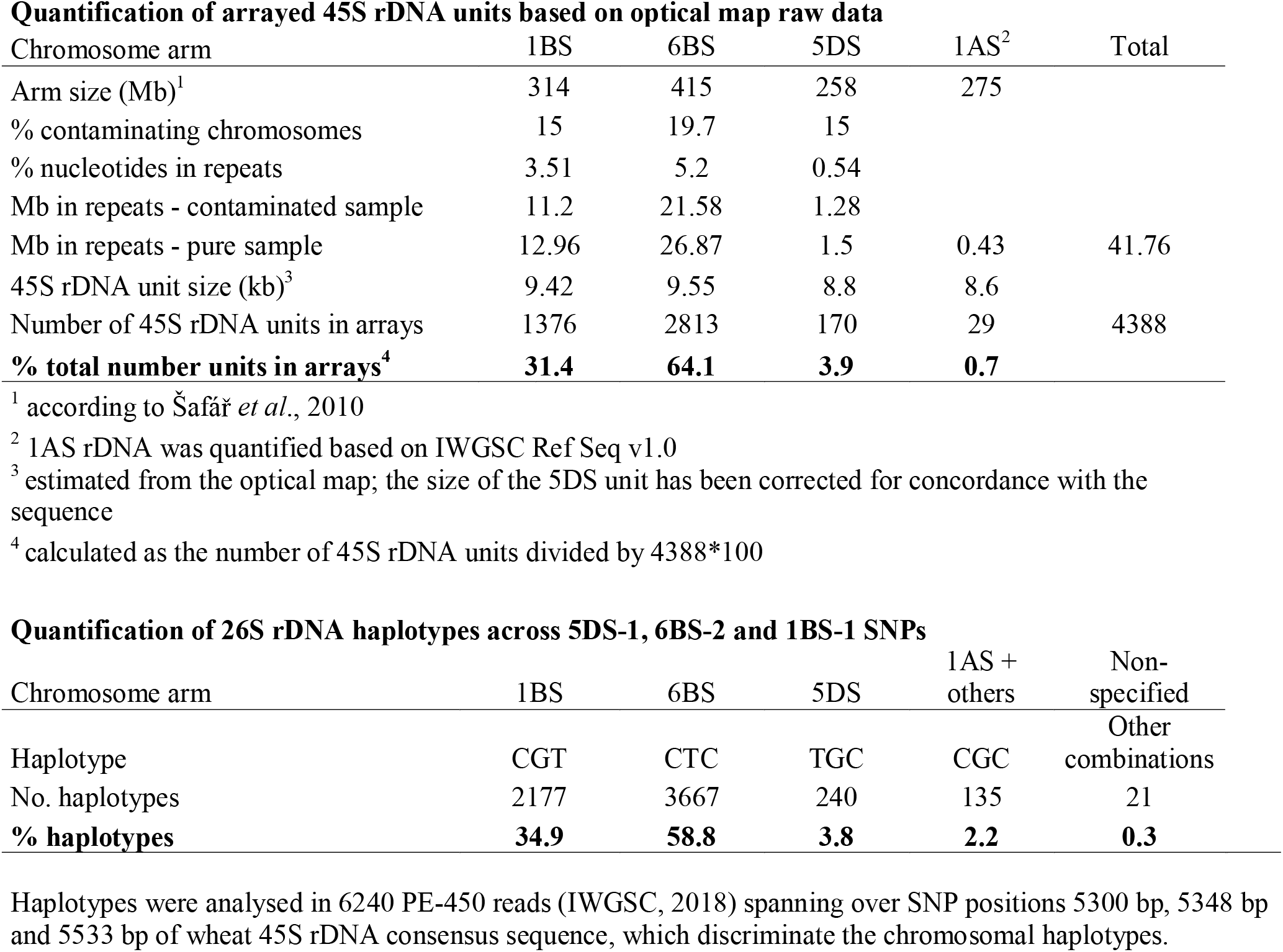
Quantification of 45S rDNA units

### Reconstruction of wheat rDNA units

A wheat consensus and three chromosome-specific 45S rDNA units were reconstructed from published short-read data using RepeatExplorer pipeline (Novák *et al*., 2013), as previously described in Kapustová *et al*. (2019). The consensus sequence was generated from whole-genome Illumina data of wheat Synthetic W7984 (SRP037990, Chapman *et al*., 2015) while the chromosome-specific units were reconstructed from Illumina reads for CS 1BS (ERX250504, IWGSC 2014) and Roche/454 reads for CS 6BS (DRX007672, Tanaka *et al*., 2014) and 5DS arms (ERA296180, Akpinar *et al*., 2015). For the unit reconstructions, a random data set of 4,471,956 filtered reads, corresponding to 0.13x genome coverage, was used for the wheat rDNA consensus, and reads corresponding to 0.15-0.77x coverages of particular chromosome arms were applied for the chromosomal consensuses. In case a chromosomal rDNA unit was not reconstructed to a contiguous sequence after one round of a graph-based clustering, the available contigs were used to extract all putative reads homologous to 45S rDNA using BLASTn. A second round of clustering was done on a data set enriched for putative rDNA reads and lacking other abundant repeat types. Finally, particular components of the 45S rDNA unit (genes and spacers) were ordered and annotated according to a rye 45S rDNA (JF489233.1, Fluch *et al*., 2012), except transcription start site, which was determined according to Vincentz and Flavell (1989).

### Identification and validation of diagnostic SNPs

Chromosome-arm-specific SNPs in coding regions were identified after aligning consensual rDNA sequences of 1BS, 6BS and 5DS in Geneious v7.1.2 (http://www.geneious.com). Diagnostic value of all putative SNPs was tested by cross-mapping of available telosome-derived Illumina reads (IWGSC, 2014) from 1AS (ERX250501), 1BS (ERX250504), 6BS (DRR008486) and 5DS (ERX250533) to individual chromosomal rDNA consensuses. Representation of particular SNPs in three *T. aestivum* ‘Chinese Spring’ assemblies (SRX2994097, Zimin *et al*., 2017; ERX1700146, Clavijo *et al*., 2017; SRX3059308, IWGSC, 2018) was assayed by mapping whole-genome Illumina reads generated in particular projects to the wheat consensual rDNA unit. The reads were filtered for quality and length and aligned to wheat rDNA consensus sequence using Bowtie2 aligner with default parameters (Langmead and Salzberg, 2012). The resulting SAM files were converted to BAM files, sorted by SAMtools package version 1.6 and 0.1.18, respectively and viewed in Integrative Genomics Viewer v2.7.2, which was also used to quantify particular SNP variants. To verify the proposed locus-specific haplotypes, we exploited merged 2 x 250 bp (PE450) reads from whole-genome libraries generated for the IWGSC RefSeq v1.0 genome (SRP114784; IWGSC, 2018) and aligned them to the wheat rDNA consensus. Reads overlapping all three diagnostic SNPs were extracted by SAMtools and checked for haplotype constitution using JVarkit Biostar214299 utility (Lindenbaum, 2015).

### RNA-seq

Five plant tissues -embryo, coleoptile (including plumule), root tips, primary leaf and mature leaf – each in two biological replicates were used for transcriptome analysis. Seeds were incubated for 5 days at 4°C in Petri dishes on wet cellulose covered by filter paper, followed by incubation at 26°C in the dark for 4-6 hours for embryo and 24 hours for coleoptile and root tips. After 24-hour incubation at 26°C, five germinating seeds were planted in garden soil and maintained in a growth chamber under long-day (16 hours’ day light) conditions and 20°/16°C for five days or until the flag leaf was visible (Zadoks’ stage 37) to collect the primary and the mature leaf, respectively. Three embryos, 2-5 coleoptiles, 10 root tips about 2mm in length and up to 100 mg of the primary leaf tissue and the mature leaf blade, respectively, were collected per sample and flash-frozen in liquid nitrogen. Total RNA was isolated using RNEasy Plant Mini Kit (Qiagen, Germany) and RNA quality was checked on Bioanalyzer using Agilent RNA 6000 Pico Kit (Agilent Technologies, Santa Clara, USA). Sequencing libraries were prepared by NEBNext® Ultra™ II Directional RNA Library Prep Kit for Illumina (New England Biolabs, Ipswich, USA) and pair-end sequenced on the Illumina NextSeq550 System. Summary of the sequencing output is provided in Table S2 **[Supplementary Information]**.

### Calling of transcript variants

Raw RNA-seq data were quality controlled using fastqc, trimmed with trimgalore/0.6.2 using default settings and rid of chloroplast and mitochondrial DNA by mapping to the respective references (accession number KC912694.1, Middleton *et al*., 2014; accession number MH051716.1, IWGSC, 2018) using HiSat2 v2.1.0 (Table S2)**[Supplementary Information]**. Two million (one million of pairs) of clean reads were used as an input for SNV calling. Out of these 90-96 % (Table S2) **[Supplementary Information]** were mapped to the wheat rDNA consensus unit, which included the external transcribed spacer (ETS)-18S-ITS1-5.8S-ITS2-26S subregions (Fig. S1A) **[Supplementary Information]**. Mapping was carried out using commands in the CLC genomics workbench (Qiagen, Germany) with the following parameters: Match score – 1, mismatch cost – 2, insertion cost – 3, deletion cost – 3, length fraction – 0.5, similarity fraction – 0.8. Read tracks were visually checked in the program window and coverage graphs were constructed. Variants were called via the ‘Probabilistic Variant Detection’ function tool in CLC using default settings. SNPs were filtered as follows: minimum read coverage – 400, count (the number of countable reads supporting the allele) −40, frequency (the ratio of “the number of ‘countable’ reads supporting the allele” to “the number of ‘countable’ reads covering the position of the variant”): ≥10% (high-frequency SNPs). Frequency of rRNA variants across five tissues were evaluated by a Pearson’s chi-squared test.

### Mapping of Iso-Seq data

To identify possible rare transcripts originating from 5DS and other minor rDNA loci, available wheat CS Iso-Seq data from PRJEB15048 (Clavijo *et al*., 2017) were mapped to the wheat rDNA consensus and 5DS rDNA unit, respectively. A total of 817,892 high-quality (CCS) Iso-Seq reads, divided into six sets corresponding to six plant developmental stages -leaf, root, seed, seedling, stem and spike -were mapped using minimap2 software with default parameters. Resulting SAM files were converted into BAM files and sorted using SAMtools. The data were visualised and variant proportion was estimated using Integrative Genomics Viewer (IGV).

### Quantification of rRNA

A hybrid reference wheat genome was generated by concatenating repeat-and rDNA-masked IWGSC RefSeq v1.0 genome sequence, the wheat consensus 45S rDNA unit, and the wheat chloroplast (KC912694.1) and mitochondrial (MH051716.1) genomes. Trimmed RNA-seq data were mapped to this reference using HiSat2 v2.1.0 (-k 1), followed by running the featureCounts function from the package Subread v1.5.2 in a paired-end mode using ‘iwgsc_refseqv1.0_HighConf_wheat_rDNA.gtf’ annotation file (IWGSC, 2018). Per-genome counts were obtained using counts and lengths of corresponding wheat nuclear (18S, 26S), chloroplast (16S, 23S) and mitochondrial (18S, 26S) rRNA genes. For mitochondria, comprising three and two non-clustered copies of 18S and 26S rRNA genes, respectively, counts for each of the genes were summed. Other rRNA types were not considered. For relative quantification of the nuclear/chloroplast/mitochondrial 18/16S and 26/23S rRNA gene transcripts, TPM (Transcripts Per Kilobase Million) values were calculated that normalize for both gene length and sequencing depth.

### Methylation of rDNA

To evaluate the level of DNA methylation in particular rDNA loci, an artificial reference sequence was compiled of the chromosomal rDNA units specific for 1BS, 6BS and 5DS (Dataset S1) **[Supplementary Information]**, respectively, which were tailed with the initial 1000 bp of the IGS to mimic concatenation. The reference was used to map three replicas of published wheat bisulphite-sequencing (BS-seq) datasets (SRR6792673, SRR6792679, SRR6792686, SRR6792688, SRR6792689, SRR6792681, SRR6792684, SRR6792687, SRR6792678, IWGSC 2018), obtained from two-week-old CS leaf tissue. The raw sequences were trimmed from the adaptors and low-quality reads were removed using Trim Galore! v0.6.2 (trim_galore -j 4 --paired --trim1). Subsequent analysis was run in Bismark-0.22.1 program (Krueger and Andrews, 2011), making use of SNPs specific for particular chromosomal loci (Dataset S2) **[Supplementary Information]**. Mapping by Bismark results in the best unique alignment by default and was run with additional parameters ‘-I 0 -X 1000 --non_directional’. The mapping was followed by deduplicate_bismark and bismark_methylation_extractor ‘--paired-end --comprehensive --no-overlap --CX –bedGraph --counts’. Methylation level in CpG, CHG and CHH contexts was calculated as a percentage of methylated cytosines (mC) out of total cytosines (C+mC) per region −5’ETS or selected part of the 26S rRNA gene) -using SeqMonk. Differential methylation between the three chromosomal units was statistically evaluated using ANOVA test and using two-tailed t-test for all pairs of chromosomal units and methylation contexts.

## RESULTS

### Positioning and characterization of rDNA arrays

Using bread wheat whole-genome Illumina reads and RepeatExplorer pipeline, we reconstructed a wheat consensus 45S rDNA unit with the length of 8193 bp (Dataset S1) **[Supplementary Information]**. We searched the unit for GCTCTTC and CTTAAG motifs, which are the recognition sites of Nt.BspQI and DLE-1 enzymes, respectively, used to label DNA molecules on optical mapping platforms of Bionano Genomics. The rDNA unit comprises one CTTAAG and two GCTCTTC sites, all of them located in the 26S rRNA gene. Since the two GCTCTTC motifs were positioned just 1126 bp apart, they could not be spatially discriminated and were expected to be recorded as a single label. Thus, each of the enzymes was predicted to introduce one label per unit and an array of tandemly organized rDNA units was expected to generate a regular labelling pattern with a spacing of 8-9 kb. To verify this prediction and to visualize particular rDNA arrays in their genomic context, we constructed chromosome-arm-specific Bionano optical maps for wheat 1BS, 6BS, 5DS and 1AS chromosome arms, respectively, which bear rRNA multigene loci. Parameters of the chromosomal maps are given in Table S1 **[Supplementary Information]**. The optical map contigs were aligned to the IWGSC RefSeq v1.0. Among 369 OM contigs from the 1BS arm, we found several comprising a regular label pattern with the spacing of 8.8-10 kb. Only 1BS contig 447, having a longer segment with an irregular pattern, could be hereby aligned to the 1BS pseudomolecule at ∼80.7 Mb position (Fig. 1B). A non-aligned 200-kb part of this contig had a regular pattern with 9.4-kb label spacing that overhung towards the telomere, indicating a presence of an rDNA array at this position. This presumption was supported by the presence of a scaffold boundary at 80.65 Mb and of a cluster of incomplete rDNA units between 80.65-80.88 Mb. Another cluster of rDNA fragments was found at 111.1-111.2 Mb, but this region was spanned by an OM contig without a regular pattern, excluding the presence of a regular rDNA array at this position.

We analogously assigned rDNA arrays to positions 111.58 Mb and 19.01 Mb of 6BS and 5DS pseudomolecules, respectively. In both the 6BS and the 5DS, we found rDNA-comprising OM contigs at both the distal and the proximal sides of the arrays (Fig. 1B). The 5DS optical map indicated that 5DS scaffold 34514 (5D:19005535..19736721 bp), flanking the rDNA array at the proximal side, was misoriented in the RefSeqv1.0 pseudomolecule, artificially breaking the rDNA array into two parts. Clusters of incomplete rDNA units flanking the rRNA multigene loci indicated that 1BS and 6BS rRNA genes were oriented from the centromere to the telomere while the 5DS rRNA locus had the opposite orientation. The rDNA-bearing OM contigs showed a highly regular pattern in 1BS and 5DS but the regularity has been disrupted in several positions by label-free gaps of tens of kilobases in the 6BS, suggesting that the rDNA array has been invaded by larger blocks of transposable elements or interspersed by shorter arrays of unlabelled tandem repeats.

We also searched for the characteristic pattern in OMs from the 1AS arm but failed in finding a single OM contig with a clear ∼9-kb pattern. Only 1AS OM contig 42 comprised segments resembling the rDNA pattern with more condensed spacing (around 8.6 kb) and some irregularities (Fig. 1B). This contig aligned with high confidence to the start of the 1A pseudomolecule, thus confirming the correctness of the sequence assembly in this region. In the sequence, we found 29 units comprising all of 18S, 5.8S and 26S rRNA genes, featured by high sequence and length diversity and variable orientation, and numerous rDNA fragments (Fig. 2A, C). This degenerated rDNA array spanned over 430 kb. The rDNA units were interspersed by telomeric repeats (Fig. 2B), indicating a dynamic character of the region and a loss of function. Majority of molecules involved in the assembly of the distal part of the OM contig 42 terminated with label-free overhangs, suggesting the possible presence of a regular telomeric sequence in the immediate vicinity of the rDNA array.

**Fig 2.**
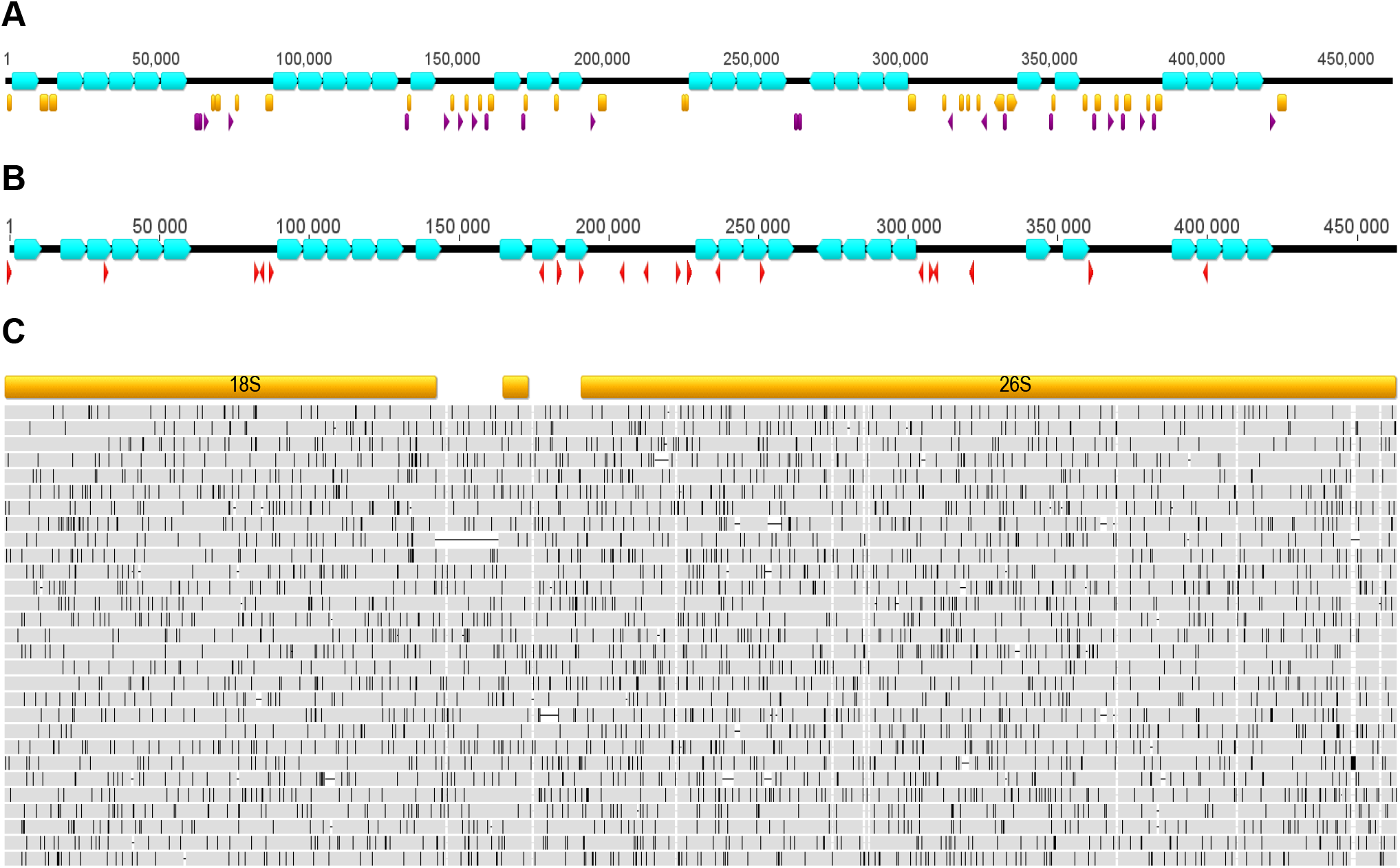
Degenerated 45S rDNA array in wheat chromosome arm 1AS. (A) The 1AS rDNA array is composed of both complete (comprising all of 18S, 5.8S and 26S rRNA genes; blue boxes), and incomplete units. Separated 18S and 26S rRNA genes and gene fragments are marked purple and yellow, respectively. (B) Ribosomal DNA units are interspersed by telomeric repeats (red triangles) of various orientation. (C) Comparison of genic parts of 29 complete 45S rDNA units building the 1AS array shows a high level of diversity. Grey bars represent individual rDNA units, disagreements are indicated by black lines.

### Quantification of rDNA units in arrays

The optical map contigs comprised segments with rDNA-specific patterns spanning over 200-370 kb, but it was obvious that these lengths did not correspond to the full size of the rDNA arrays, but rather reflected the sizes of DNA molecules used for optical map assembly, which did not exceed 500 kb. Still, an approximate quantification could be done by the IrysView software, which includes algorithms that calculate the proportion of nucleotides comprised in tandemly organised repeats of a particular size in OM raw data. Results of this analysis revealed repeats of 8.5-10 kb as the major labelled tandem repeats in chromosome arms 1BS, 6BS and 5DS (Fig. S2) **[Supplementary Information]**. We quantified tandem-organized rDNA units in each rDNA locus, employing information on the percentage of nucleotides in repeats, sizes of the telosomic chromosomes and purities of particular sorted fractions (Table 1). This analysis indicated that the largest array of rDNA units was located in the 6BS chromosome arm (2813 units, 26.87 Mb), followed by arrays in 1BS (1376 units, 12.96 Mb), 5DS (170 units, 1.5 Mb) and 1AS (29 complete units, 0.43 Mb), being the smallest locus. The total of 4388 units occupied approximately 42 Mb of the wheat CS genome space. Relative quantification of the units assigned 31.4 and 64.1% of them to the major NORs in 1BS and 6BS, respectively. The remaining 3.9 and 0.7% belong to the minor loci in 5DS and 1AS, respectively.

### Chromosome-specific rDNA units and diagnostic SNPs

Chromosome-specific consensual units were reconstructed for 1BS, 6BS and 5DS loci, featured by regular rDNA arrays (Dataset S1) **[Supplementary Information]**. Mutual comparison of the chromosomal consensual sequences (Dataset S2) **[Supplementary Information]** revealed a high sequence diversity in IGS, including 5’ETS, and several SNPs in both ITS1 and ITS2. While the 18S and 5.8S rRNA gene sequences were identical among the three chromosomal consensuses, a total of eight putative arm-specific SNPs were identified in the 26S rRNA gene: two 6BS-specific, two 1BS-specific and four 5DS-specific. Diagnostic value of all these SNPs was tested by cross-mapping Illumina reads from 1AS, 1BS, 6BS and 5DS to the three consensuses. This assay led to the selection of two diagnostic SNPs for each rRNA multigene locus. The 5DS-2 SNP variant was also found in 1A and other minor rDNA clusters outside the B genome (Table S3) **[Supplementary Information]**, thus we called it “non-B”. Further, we analysed the frequency of allelic variants of the proposed diagnostic SNPs in the genome of CS wheat by investigating raw data that had been used for RefSeq v1.0 assembly (IWGSC, 2018), and also Illumina reads used for two additional CS genome assemblies (Triticum 3.1, Zimin *et al*., 2017; TGACv1, Clavijo *et al*., 2017) (Table S4) **[Supplementary Information]**. Pairs of SNPs featuring 1BS and 6BS, respectively, had highly consistent representations in IWGSC RefSeq v1.0 and Triticum 3.1 data, while showing slight inconsistencies in TGACv1, possibly due to a small number of available reads or difference in CS accessions used in particular projects. The RefSeq v1.0 analysis showed that the 1BS, 6BS and 5DS repeat unit variants represented 36.2, 56.6 and 4.2% of the whole-genome data, respectively. Similar representation of particular chromosomal loci has been found in raw data of the other CS projects.

Based on this analysis, we proposed locus-specific haplotypes and verified them using merged 450-bp Illumina pair-end reads, applied for the IWGSC RefSeq v1.0 assembly. Alignment of the reads to the wheat consensus rDNA (Fig. S3) **[Supplementary Information]** showed 6240 of them covering a triplet of chromosome-specific SNPs, 5DS-1, 6BS-2 and 1BS-1, that enables discrimination of haplotypes proposed for 1BS, 6BS and 5DS loci, respectively. Out of the 6240 reads, 2177 (34.9%) corresponded to the predicted allelic constitution of 1BS, 3667 (58.8%) to 6BS and 240 (3.8%) to 5DS. Another haplotype, observed in 135 (2.2%) reads was found in most units of the 1AS array and in small clusters of degenerated rDNA units in 7DS and other arms, which besides that comprised units with the 1BS haplotype. Only 21 (0.3%) 450-bp reads had a different combination of SNPs (Table 1). The results of the haplotype analysis thus supported validity of the proposed diagnostic SNPs.

### Locus-and tissue-specific transcription of 26S rDNA

To assess locus-specific transcription, we used short Illumina RNA-seq and long Iso-Seq (PacBio) reads of high and low sequence coverage, respectively. A typical profile of RNA-seq read coverage along the rDNA unit is shown in Fig. S1B **[Supplementary Information]**. As expected, the coverage was high in genic regions and low in the non-coding regions. Depending on the tissue, only 0.006-0.122% reads mapping to wheat consensus rDNA unit belonged to the 5’ETS, that represents 16.3% of the primary 45S rDNA transcript length. This implies that just a fraction of a percent of sequenced rRNAs had full-length 5’ETS, indicating that a minute amount of 45S pre-rRNA was present in the sequenced RNA samples. The highest and the lowest 5’ETS coverage was observed in the embryo and the primary leaf, respectively.

The analysis of high-frequency (>10%) variants in the RNA-seq data obtained from five tissues confirmed the presence of significant transcript variants at the diagnostic SNPs in both major rDNA loci in the B genome, namely 6BS-1, 6BS-2, 1BS-1 and 1BS-2. In all tissues, the 1BS rRNA variants predominated those of the 6BS in a ratio roughly 2:1, which slightly varied among tissues (Fig. 3A; Table S5) **[Supplementary Information]**. The highest frequency of 6BS-specific variants was in the coleoptile (35 and 37.6% in 6BS-1 and 6BS-2, respectively) while the highest frequency of the 1BS-specific variants was in the mature leaf (71.9 and 75.1% in 1BS-1 and 1BS-2, respectively).

**Fig 3.**
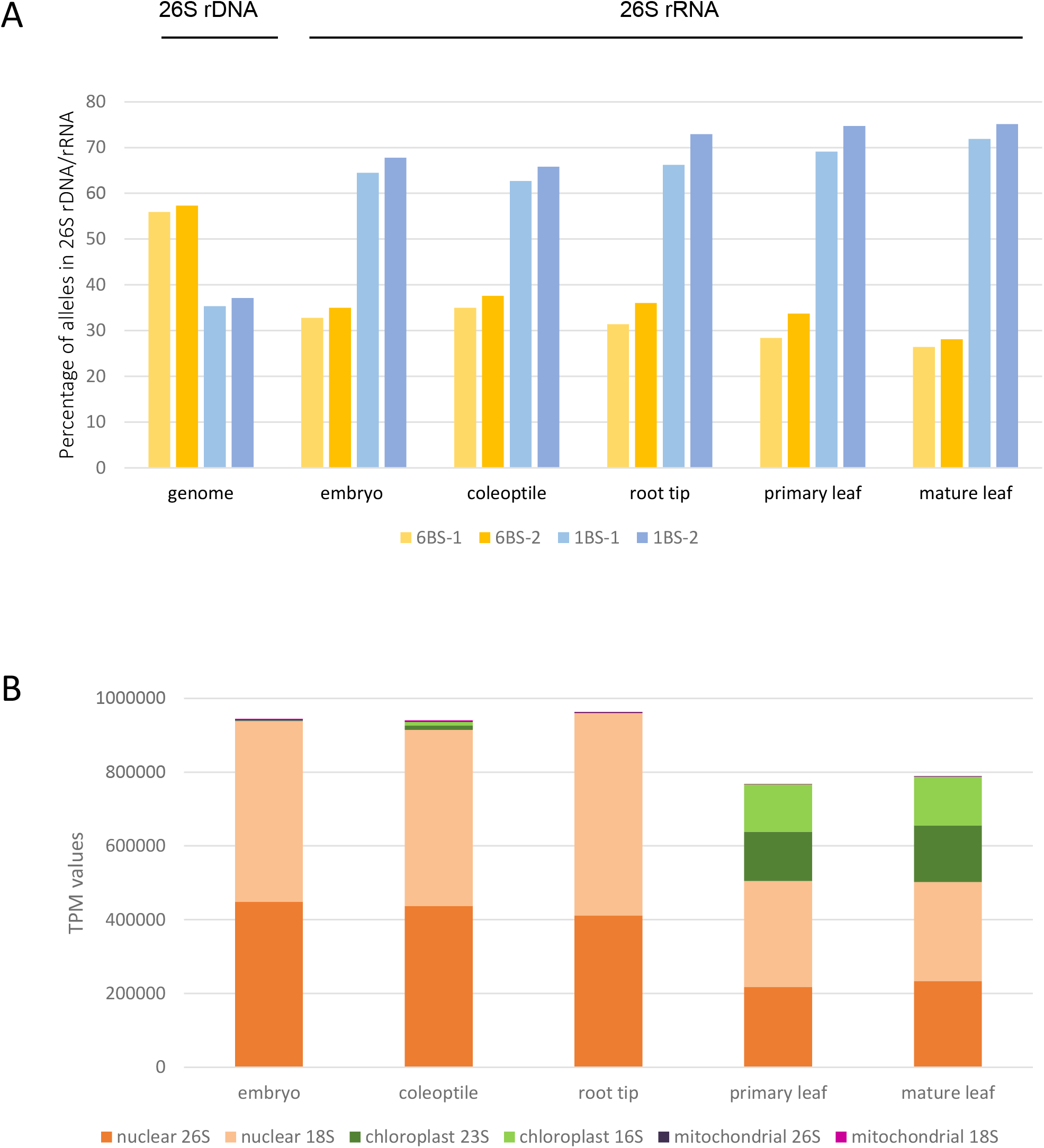
Tissue-specific contribution of particular loci to rRNA production. (A) Representation of 6BS-and 1BS-specific SNP variants in 26S rDNA, as estimated from raw Illumina data used to generate IWGSC RefSeq v1.0 (*genome*), and in 26S rRNA from five tissues (*embryo, coleoptile, root tip, primary leaf, mature leaf*). (B) Relative contribution of nuclear, mitochondrial and chloroplast genomes to overall rRNA production across five tissues, expressed as TPM (Transcripts Per Kilobase Million) values for each of the 16S/18S and 23/26S rRNA types. The graphs show mean values of two replicas.

Our failure to identify allelic variants in 5DS-1 and 5DS-2 SNP positions suggested the absence of transcription outside the major loci. To validate this finding, we inspected publically available Iso-Seq data of CS wheat with reads of several kilobases in length, promising to reliably identify possible 5DS transcripts. The majority of transcripts in all datasets started at position 2351 of the rDNA consensus, which is the start of the 18S rRNA gene, and did not extend beyond position 8154 (Fig. S4A, B) **[Supplementary Information]**, preceding a polypyridine CCCTCCCCC tract. The Iso-Seq datasets for particular developmental stages (leaf, root, seed, seedling, stem and spike) differed in read length and amount of Iso-Seq data mapped to the 45S rDNA unit. The most beneficial showed the dataset of “Seed” comprising 1951 reads mapping to the 5DS rDNA unit, which was used to search for transcripts of 5DS and possibly other minor rRNA loci. In five out of 1079 "Seed” Iso-Seq reads mapping to the position of 5DS-1 in the 5DS rDNA unit, we could unambiguously recognize the 5DS-specific haplotype based on the 5DS-1, several 5DS-specific SNPs in ITS2 (Dataset S2) **[Supplementary Information]** and, in two cases, also 5DS-2 and 1BS-2 SNPs (Fig. S4C) **[Supplementary Information]**. Additional two reads out of 425 mapping to the terminal part of the 26S rRNA gene, covering 5DS-2 and 1BS-2 but not 5DS-1, suggested a non-B origin of the transcripts, and another three reads, covering the entire 26S rRNA gene, had a non-B non-5DS combination of SNP alleles, indicating possible 1A origin. To summarize, we observed among the “Seed” transcripts of CS wheat possible non-B variants of 26S rRNA with the frequency of around one percent.

### Unequal cytosine methylation of rDNA loci

To clarify mechanisms controlling the expression of particular rRNA multigene loci, we carried out a comparative analysis of cytosine methylation assigned to particular chromosome-specific rDNA sequences. The methylation was evaluated after mapping BS-seq reads to chromosomal consensual units of 1BS, 6BS and 5DS, respectively. This approach allowed comparing methylation ratios in non-repetitive segments of the rDNA units that showed polymorphisms between chromosomal consensual sequences. The reads covered well 5’ETSs, both ITSs and the polymorphic parts of the 26S rRNA genes. The 5’ETS region of 1141bp length contained 13 evenly distributed SNPs discriminating 1BS-and 6BS-specific reads; the ETS of the 5DS unit comprised 39 SNPs that distinguished it from those of the B-genome loci (Dataset S2) **[Supplementary Information]**. The extent of cytosine methylation was quantified for the 5’ETS and the part of the 26S rRNA gene with validated SNPs (Fig. 4, Fig. S5) **[Supplementary Information]**. In the 5’ETS, the comparative analysis revealed generally high levels of DNA methylation in the CpG contexts, reaching 92.05, 94.77 and 94.47% for 1BS, 6BS and 5DS unit, respectively. The difference between 1BS and 6BS (2.72%) was highly significant (p < 0.001). On the contrary, the frequency of cytosine methylation in the CHG and CHH contexts was markedly lower (23.28, 22.97 and 43.55% for CHG, and 1.4, 1.49 and 2.31% for CHH in 1BS, 6BS and 5DS unit, respectively) and significantly (p < 0.001) higher in 5DS compared to 1BS and 6BS loci. The difference between 1BS and 6BS in the CHG and CHH contexts was non-significant. Pattern of cytosine methylation in the analysed part of the 26S rRNA gene was highly similar to that in the 5’ETS (Fig. S5) **[Supplementary Information]**, except for a noticeable increase in methylation in the CHG contexts in both 1BS (32.2 vs. 23.3%) and 6BS (30.7 vs. 23%) units.

**Fig 4.**
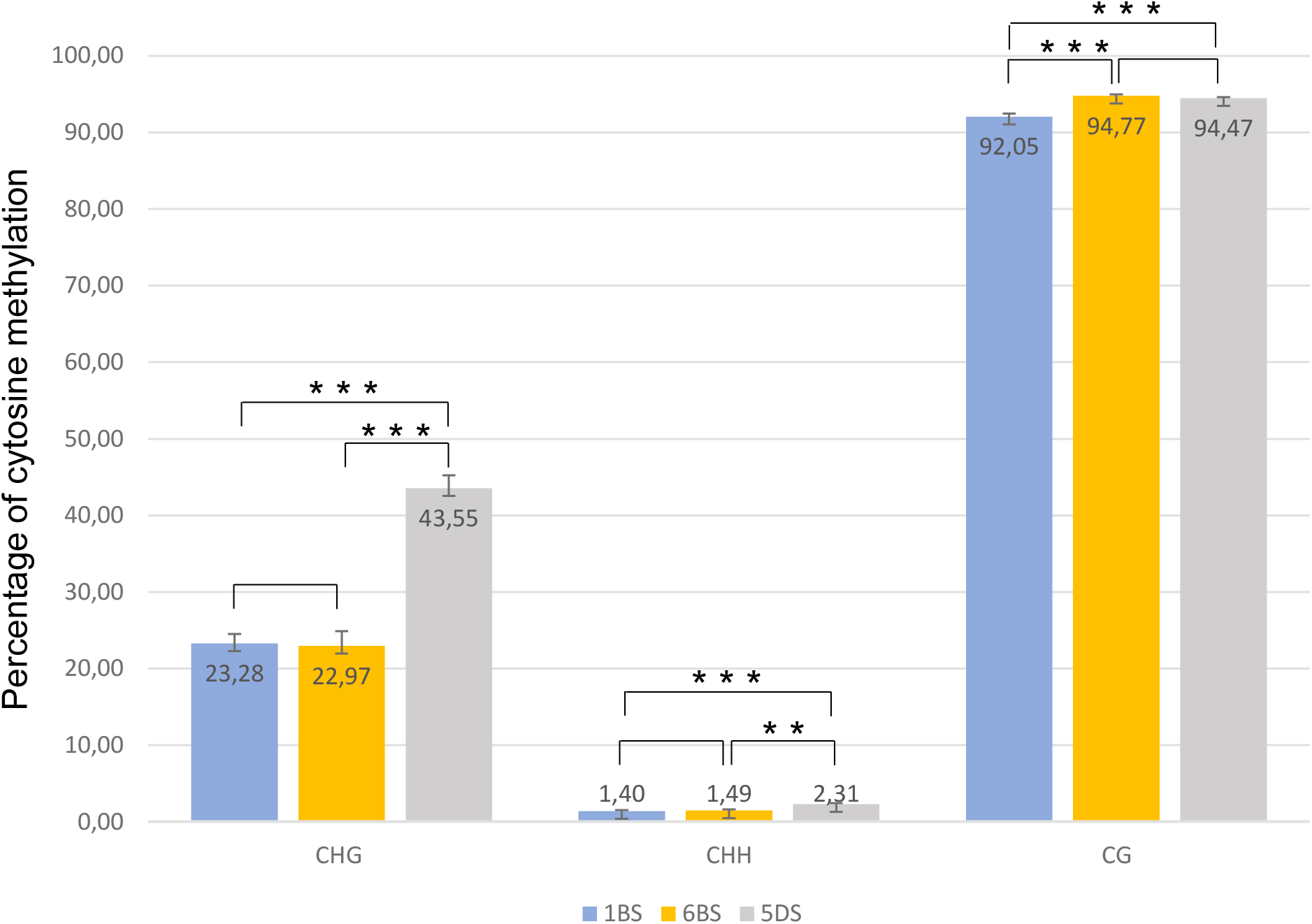
Cytosine methylation in 5’ETS of 45S rDNA unit from 1B, 6B and 5D loci. The methylation was evaluated after mapping three replicas of publically available bisulfite-sequencing reads (IWGSC, 2018) to consensual units of 1BS, 6BS and 5DS, respectively. The methylation was quantified in 5’ETS of each unit as percentage of methylated cytosine in CpG, CHG and CHH contexts, respectively. Asterisks indicate levels of statistical significance (** p < 0.01; *** p < 0.001) of differences between methylation in 1BS, 6BS and 5DS loci. Absence of asterisks indicates non-significant difference (p > 0.05).

### Dynamic expression of organellar rDNA

Relative quantification of nuclear, chloroplast and mitochondrial ribosomal RNAs across five tissues was based on TPM estimates for the large rRNAs (16/18S and 23/26S). The relative amount of rRNA in transcripts culminated in tissues with high proportion of rapidly dividing meristematic cells (embryo, coleoptile, root tip), likely reflecting a high demand on ribosome biogenesis and proteosynthesis in these tissues. As expected, production of plastid rRNA was minute in embryo, coleoptile and root tip but the proportion of chloroplast rRNA increased dramatically in the leaf samples, in line with the proliferation of chloroplasts and high photosynthetic activity. Alongside this chloroplast rRNA increase, the proportion of nuclear rRNA dropped to about one half in the leaf samples. The relative contribution of mitoribosomal RNA to overall rRNA production was negligible in all tissues (Fig. 3B; Table S6) **[Supplementary Information]**.

## DISCUSSION

### Ribosomal DNA quantity and distribution

Instability in rDNA copy number has been well documented in both plants and animals at the inter-population and even inter-individual levels (Rogers and Bendich, 1987; Rabanal *et al*., 2017, Nelson *et al*., 2019). In our study, we used a new approach to quantify 45S rDNA in the genome, which was based on calculating tandemly organized repeats of the corresponding size in optical map raw data. Our total estimate −4388 rDNA units in arrays -was remarkably similar to that of Flavell and O’Dell (1976) who, based on classical filter hybridization techniques, estimated 4575 copies per 1C genome of CS wheat. The accordance between both studies suggests robustness of both quantification approaches and points to relative stability of rDNA copy number, even between standard karyotype of CS wheat and derived telosomic lines used in our study, which have undergone multiple meiotic cycles since their generation in 1970’s (Sears and Sears, 1978). Nevertheless, a recent study of Handa *et al*. (2018) reported 11,160 rDNA units in the CS genome, which is roughly 2.5 higher estimate than that of Flavell and O’Dell (1976) and ours. The incongruence between the studies can be assigned to different approaches used to quantify the rDNA units rather than to inter-individual variability. The hybridization techniques used by Flavell and O’Dell (1976) are likely to pick up only longer DNA sequences while qPCR used by Handa and co-workers can target also short rDNA-unit fragments of a few hundred base pairs, which are abundant in marginal parts of rDNA arrays and are also dispersed across the genome (Fig. 1). The optical-mapping approach applied in the current study considered only units organized in arrays, which are likely to be functional, while qPCR cannot discriminate the arrayed genes from the dispersed ones that are frequently truncated or recombined and likely pseudogenized. Of note, BLAST search for the consensual rDNA unit in the IWGSC RefSeq v1.0 genome, carried out in Ensembl Plants (https://plants.ensembl.org/Triticum_aestivum/Tools/Blast), provided 1431 hits of high confidence (E-value < e^-100^), distributed on all pseudomolecules (743 hits) and in unassigned scaffolds (688 hits). Majority of them were located outside regular arrays and represented rDNA fragments. Taking this into account, we consider our approach more relevant for assessing sequences with potential to contribute to rRNA production. Nevertheless, we cannot exclude that our outcomes slightly underestimate the number of rDNA units due to false negative/positive labels occurring occasionally in the OM raw data and possibly disrupting the regularity of the label pattern.

The relative proportion of particular rDNA loci was determined both from the optical map data and from the representation of locus-specific haplotypes in whole-genome data (Table 1). The results obtained by both approaches were concordant, except for a slight shift (34.9 vs. 31.4%) in favour of 1B in the haplotype data. This may reflect the fact that the haplotype analysis does not exclude the non-arrayed units and larger unit fragments dispersed across the genome where the 1BS haplotype predominates, probably due to the higher transcription activity. The 1BS: 6BS ratio of 31.4: 64.1 is in a good agreement with 30/30.5: 60/60.9 obtained in previous studies (Flavell and O’Dell, 1976; Handa *et al*., 2018) but our OM data ascribe a smaller percentage to other chromosomes (4.6 vs. 10/8.6%), for the reasons explained above. The occurrence of haplotypes in 26S rRNA gene with high locus specificity suggests limited interchromosomal conversion events between particular arrays, which is supported by observations of Lassner *et al*. (1987) who did not find evidence of homogenization between B-and D-genome rDNA based on comparing IGS sequences from the 6B and 5D loci. On the other hand, Handa *et al*. (2018), analysing 3’ETS region in 45S rDNA, identified two rDNA subtypes that were shared between 1B and 6B chromosomes, suggesting sequence interchange between the two chromosomes.

### Nucleolar dominance of B-genome NORs

To assess the transcription of particular chromosomal rDNA loci, we analysed data obtained by high-throughput sequencing of total RNA, making use of validated diagnostic SNPs located in 26S rRNA gene, which was well covered by transcripts. The results for 1B-and 6B-specific transcription supported findings of Martini and Flavell (1985) who measured volumes of nucleoli belonging to *NorB1* and *NorB2*, respectively, and predicted 2:1 transcription ratio for the 1B and 6B loci. Fractional transcription from the 5DS locus detected in our study is in agreement with observations of Mukai *et al*. (1991) who found 5DS-associated micronucleoli in ∼3% of CS nuclei. Several studies (Cermeño *et al*., 1974; Martini and Flavell, 1985; Handa *et al*., 2018) observed activation of minor rDNA loci, mainly *Nor-D3* in 5D, after removal of one of the major NORs. This is consistent with our finding that the 5DS locus is represented by a regular array of compactly organized rDNA units, suggesting their full functionality. The silenced state of the 5DS locus is likely maintained by epigenetic mechanisms, among which cytosine methylation in CHG and possibly CHH contexts appears to play a significant role, as suggested by our comparative analysis of DNA methylation levels in 1BS, 6BS and 5DS-specific 45S rDNA units (Fig. 4, Fig. S5) **[Supplementary Information]**. Similarly, in a study of Guo and Han (2014), promoter methylation in CHG and CHH rather than CpG contexts was involved in silencing of A-genome rDNA loci in synthetic allotetraploid wheats. Given that the CHG methylation is ultimately associated with heterochromatic H3K9m2 histone mark (Lindroth *et al*., 2004), it is likely that the 5D NOR is in phase of heterochromatization and permanent inactivation.

We failed to find clear evidence for 26S rDNA expression from the 1AS locus that is featured by large scale rearrangements and truncated copies, indicating extensive pseudogenization, which suggests that silencing of this locus is most probably terminal (mutation-directed), in contrast to that of 5DS. Still, several non-truncated units seem to be present in 1AS, harbouring shorter IGS than in other loci. Given that lengths of the IGS were correlated with transcription activity in wheat (Sardana *et al*., 1993), it is likely that the 1AS array became underdominant early after hybridization of the A and B genomes, which gave rise to tetraploid AABB wheat, and that the rearrangements arrived later in the evolution. This is supported by the observation of Guo and Han (2014) that the A genome loci were silenced in the synthetic lines of wheat and eventually lost in subsequent generations. That study also demonstrated nucleolar dominance of the D over the A and the B over the D genome in newly formed allotetraploids. Interestingly, our evidence of an irregular rDNA array in 1A and a potentially functional array in 5D, whose suppression was reverted in several studies, indicates that the process of elimination of the subdominant rDNA loci has been much slower in natural bread wheat than in the synthetic polyploids reported by Guo and Han (2014). Since the D genome has been a relatively recent addition to the hexaploid bread wheat genome (IWGSC, 2014), the difference in the extent of suppression between the two minor loci may reflect the length of co-existence of the A and the D with the dominant B genome. Alternatively, the lower suppression of the D genome might be due to higher competitiveness of its rDNA, possibly due to longer IGS compared to the A genome.

### Uneven transcription of the major B-genome rDNA loci

In contrast to the B genome nucleolar dominance, explored in several studies, the partial predominance of 1B over 6B rDNA transcription gained less attention and its mechanism has not been clarified yet. Handa *et al*. (2018) suggested a regulation mechanism based on rDNA sequence difference because their predominantly expressed subtype of 3’ETS sequence was found on both 1B and 6B chromosomes. In our study, and also in that of Martini and Flavell (1985), the higher expression was associated with the NOR in 1B chromosome. Similarly, the transcription activity was found associated with a particular chromosome in a diploid species *Arabidopsis thaliana*, (Chandrasekhara *et al*., 2016). Since the active and the developmentally silenced NORs, located in *A. thaliana* chromosomes 4 and 2, respectively, are comparable in size, unit sequence and position on the chromosome in the Col-0 ecotype, the authors proposed a critical role of a *NOR2*-adjacent region in initiating the selective silencing of the locus. However, the mechanism of spreading heterochromatization appears unlikely in the partial silencing of the 6B locus since wheat NORs were found to form nucleoli with expressed (decondensed) and unexpressed (condensed) rDNA segments interspersed, in contrast to other species (Leitch *et al*., 1992).

Obviously, the expression dominance of the 1B NOR is rather labile and could be genotype-specific and unit-structure-dependent. For example, another wheat variety, *T. aestivum* ‘Bezostaya’, having larger 6B rDNA units, expressed the 6B NOR more efficiently than that in 1B, as concluded from cytogenetic observations (Sardana *et al*., 1993). The authors suggested that the IGS size, specifically the number of 135-bp repeats therein, and methylation status of a specific CCGG site downstream of the repeats might be decisive for transcription activity of particular rDNA loci in wheat. They assumed for 1B units in CS a larger IGS size compared to those in 6B and 5D, as demonstrated in several studies (Appels and Dvořák, 1982; Lassner *et al*. 1987; Flavell *et al*. 1988). Our optical map data (Table 1) confirm a larger rDNA unit size for 1B vs. 5D but not 6B locus, whose unit mean size appears ∼130 bp larger than that of the 1B locus. Nevertheless, the analysis of OM single-molecule data revealed for each chromosomal locus a broader range of unit sizes (Fig. S2) **[Supplementary Information]**, suggesting that rDNA arrays in particular chromosomal loci are not completely homogenous, at least in terms of the unit size. Consistent with this, Flavell *et al*. (1988) found the longer spacers only in a small fraction of rRNA genes from the 1B NOR of CS. Such heterogeneity might eventually result into a variability in the proportions of longer (possibly highly active) and shorter (moderately active or inactive) units, which would explain observation of Mukai *et al*. (1991) who found the two major loci equally active in CS, based on cytogenetic analyses.

### Developmental control of rRNA dosage

We carried out transcriptomic analysis across five wheat tissues aiming to identify differential transcription of 45S rDNA, as observed in both diploid and allopolyploid species of other genera (Chen and Pikaard, 1997; Komarova *et al*., 2004; Pontvianne *et al*., 2010; Dobešová *et al*., 2015). Here we observed moderate developmental silencing of the 6B rDNA locus, which was most apparent in the adult leaf tissue. The higher expression of the 6B-specific rDNA in early-stage tissues with rapidly dividing cells (embryo, coleoptile, root tip), might be conducted by their overall high demands on nuclear rRNA production, as suggested by our transcript analysis across tissues (Fig. 3B). Studies in *A. thaliana* ascribed the major role in the developmental regulation of minor variants in both 45S and 5S rDNA to chromatin state of the loci, related to the proportion of meristematic cells in particular tissues (Mathieu *et al*., 2003; Pontvianne *et al*., 2010). The expression of a developmentally silenced 45S rDNA variant *VAR1*, specific for the *NOR2*, was found associated with a lower degree of chromatin condensation in the early stage of seedling development and also with decreased cytosine methylation in 5’ETS of 45S rDNA. The *VAR1* expression was apparent up to the second day of seed germination and in later stages only weakly in root and floral tissues, while completely disappearing from leaves (Pontvianne *et al*., 2010). This expression pattern is consistent with the trend observed for the 6B locus in our transcriptomic data (Fig. 3A), thus we propose an analogous mechanism to control the developmental dynamics of rRNA transcription in wheat. Nevertheless, the extent of 6B-NOR silencing detected in our study was less significant, perhaps due to small differences in IGS sizes between 1B and 6B loci, which appear determining for the unit methylation status and transcription activity of rRNA genes in wheat and its relatives (Flavell *et al*., 1988; Sardana *et al*., 1993). We observed small, but highly significant difference in the level of CpG methylation between 1B and 6B units and no difference in CHG methylation (Fig. 4, Table S7) **[Supplementary Information]**. This subtle variance might reflect the fact that only a minor part of rDNA units in both 1B and 6B NORs appear active (Mukai *et al*., 1991), thus diminishing potential differences in global epigenetic patterns between the two loci.

### Conclusions

Our study provided comprehensive information on fine structure of major and minor rDNA loci and their activity and evolution in bread wheat genome (summarised in Fig. 5). We documented various mechanisms of rRNA dosage control, including gene elimination and stable inactivation related to nucleolar subdominance of the A and D-genome loci, and partial, developmentally regulated silencing of one of the major B-genome loci. To fully understand the latter mechanism, further studies across developmental stages will be needed to assess involvement of epigenetics and chromatin arrangement as well as the role of intergenic spacers and their transcripts.

**Fig 5.**
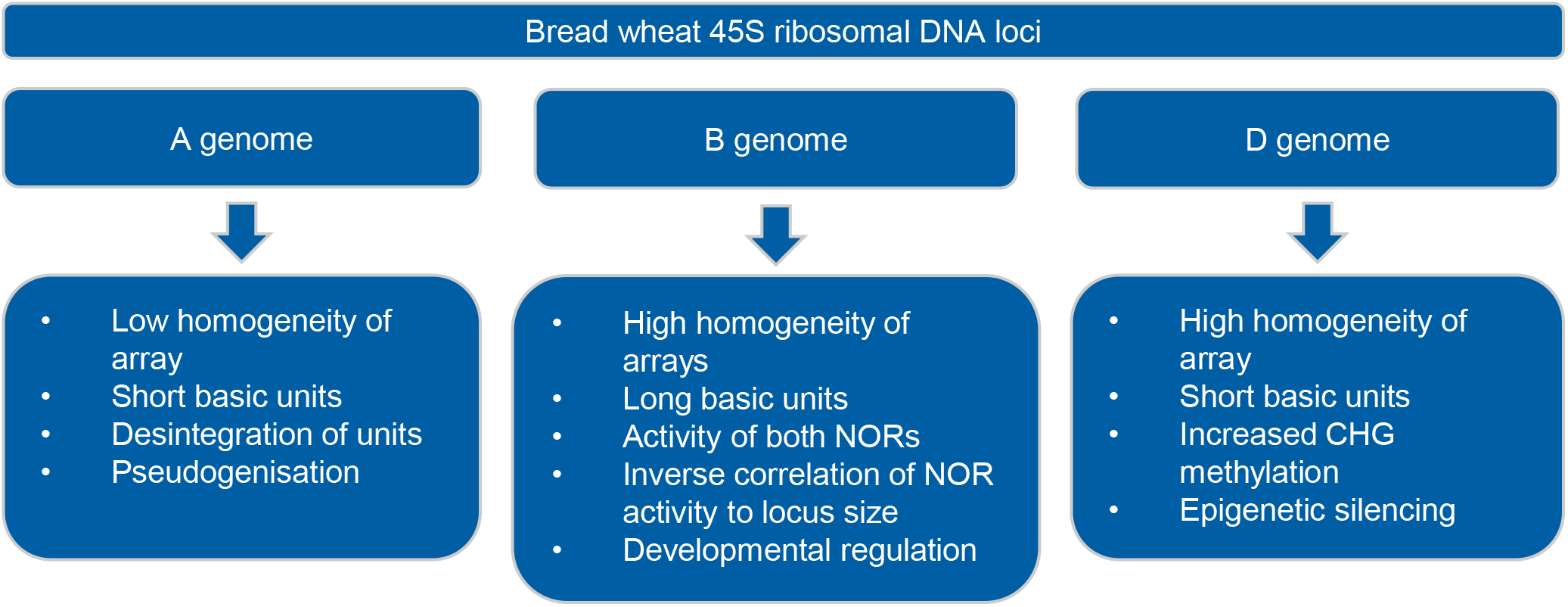
A scheme for the NOR evolution and transcription regulation in bread wheat genome

## Supplementary Data

Table S1. Optical map statistics Table S2. RNA-seq data

Table S3. Frequency of allelic variants at diagnostic SNPs in Illumina reads of flow-sorted chromosome arms

Table S4. Frequency of allelic variants at diagnostic SNPs in Illumina reads from three *T. aestivum* ‘Chinese Spring’ genome projects

Table S5. Variant analysis of 26S rRNA in different tissues -statistics

Table S6. Transcripts per kilobase million (TPM) values for nuclear, chloroplast and mitochondrial 16S/18S and 23/26S rRNAs across five tissues

Table S7. Cytosine methylation in 5’ETS of 45S rDNA units -statistics

Fig. S1. Wheat consensus 45S rDNA unit and its coverage by RNA-seq reads. Fig. S2. Quantification of 45S rDNA units in arrays from optical map data.

Fig. S3. Analysis of chromosome-specific haplotypes in 45S rDNA.

Fig. S4. Mapping of Iso-Seq reads to wheat consensus and 5DS rDNA units.

Fig. S5. Cytosine methylation in 5’ETS and 26S rRNA gene from 1B, 6B and 5D loci

Dataset S1. Sequences of wheat 45S rDNA consensus and chromosome-specific consensual rDNA units.

Dataset S2. Multiple alignment of wheat 45S rDNA consensus and chromosome-specific consensual rDNA units.

## Acknowledgements

We thank to Zdeňka Dubská, Jitka Weiserová and Eva Jahnová for excellent technical assistance. We are grateful to M. Mascher for sharing merged PE450 Illumina reads of wheat. Computational resources were supplied by the project “e-Infrastruktura CZ” (e-INFRA LM2018140) provided within the program Projects of Large Research, Development and Innovations Infrastructures, and by the ELIXIR-CZ project (LM2018131), part of the international ELIXIR infrastructure. Author contribution: HŠ planned and designed the research. AK and ZT conducted transcriptome analysis, ZT and HT carried out optical mapping, including data analysis. PN performed rRNA quantification and cytosine methylation analysis. VK validated diagnostic SNPs and together with JM a EH reconstructed consensual rDNA units. JV flow sorted chromosomes for optical mapping. HŠ, AK and JD were involved in writing the manuscript and acquisition of funding for the study.

## Data Availability

Raw RNA-seq data for five tissues generated for this study are openly available in the NCBI Short Read Archive under BioProject PRJNA657991. Consensual sequences of rDNA units and their alignments are available within Supplementary Data published online. Optical maps of 1AS, 1BS, 6BS and 5DS wheat chromosome arms (bnx and cmap files) are available from the corresponding author, Hana Šimková, upon request.

## Appendix

## Abbreviations

BS-seq: bisulphite sequencing
CS: *T. aestivum* cv. Chinese Spring
ETS: external transcribed spacer
IGS: intergenic spacer
ITS: internal transcribed spacer
IWGSC: International Wheat Genome Sequencing Consortium
NOR: nucleolus organizer region
OM: optical map
TPM: Transcripts Per Kilobase Million

