## Supplemental Figures S1-S5 for "Anatomy, transcription dynamics and evolution of wheat ribosomal RNA loci deciphered by a multi-omics approach"

### Slide 1
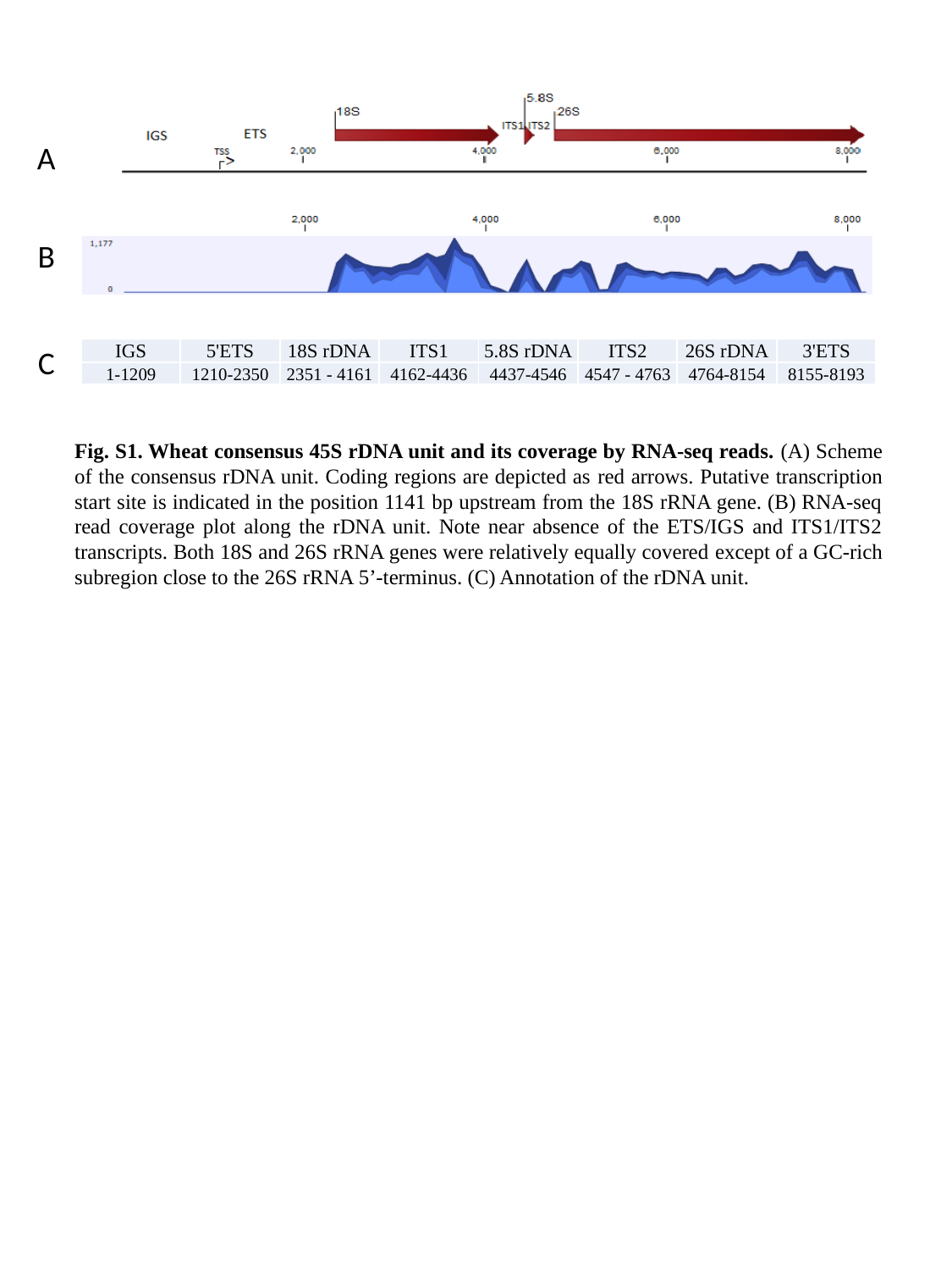

A
B
C
| IGS | 5'ETS | 18S rDNA | ITS1 | 5.8S rDNA | ITS2 | 26S rDNA | 3'ETS |
| --- | --- | --- | --- | --- | --- | --- | --- |
| 1-1209 | 1210-2350 | 2351 - 4161 | 4162-4436 | 4437-4546 | 4547 - 4763 | 4764-8154 | 8155-8193 |
Fig. S1. Wheat consensus 45S rDNA unit and its coverage by RNA-seq reads. (A) Scheme of the consensus rDNA unit. Coding regions are depicted as red arrows. Putative transcription start site is indicated in the position 1141 bp upstream from the 18S rRNA gene. (B) RNA-seq read coverage plot along the rDNA unit. Note near absence of the ETS/IGS and ITS1/ITS2 transcripts. Both 18S and 26S rRNA genes were relatively equally covered except of a GC-rich subregion close to the 26S rRNA 5’-terminus. (C) Annotation of the rDNA unit.

### Slide 2
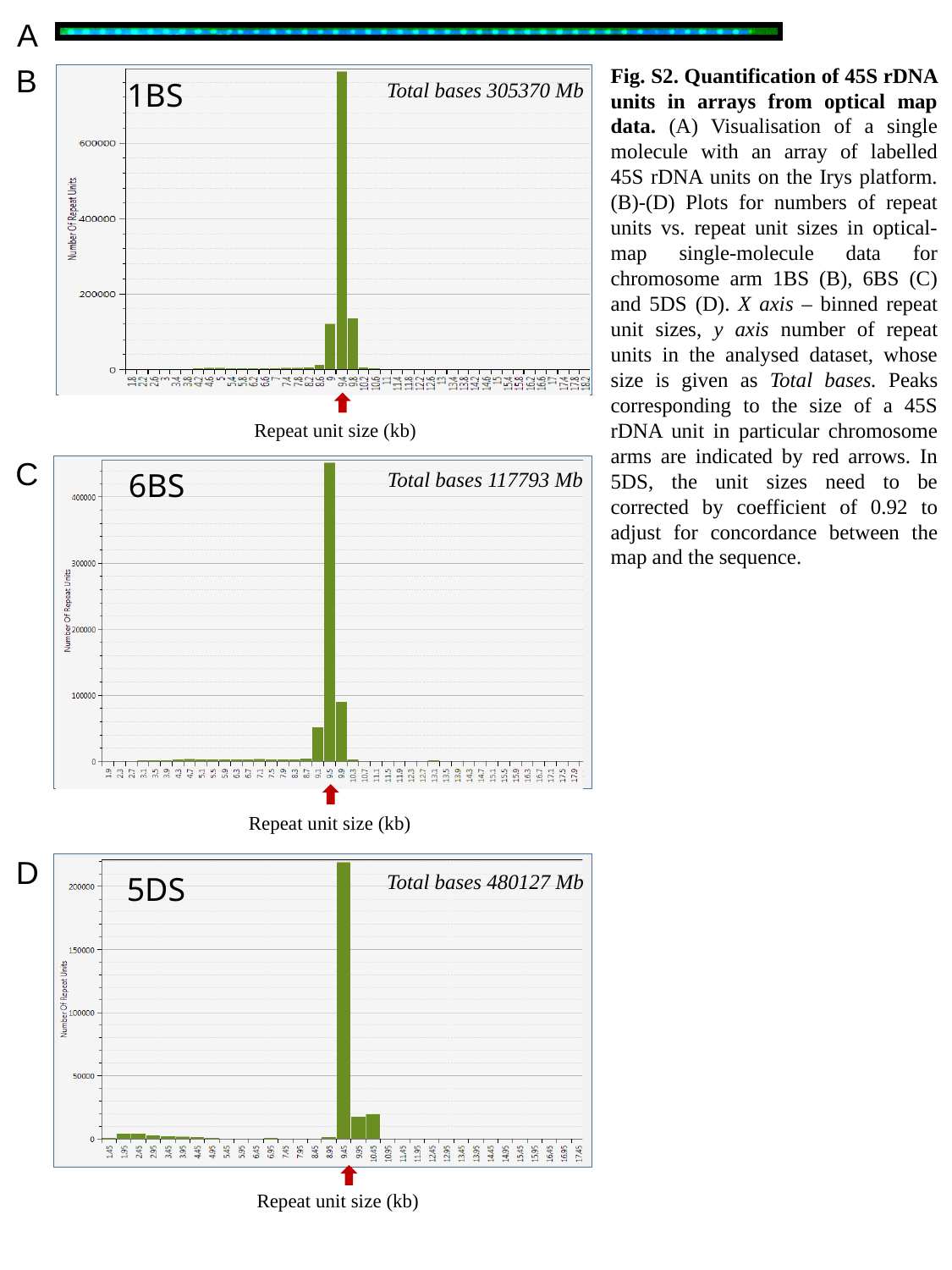

A
B
Fig. S2. Quantification of 45S rDNA units in arrays from optical map data. (A) Visualisation of a single molecule with an array of labelled 45S rDNA units on the Irys platform. (B)-(D) Plots for numbers of repeat units vs. repeat unit sizes in optical-map single-molecule data for chromosome arm 1BS (B), 6BS (C) and 5DS (D). X axis – binned repeat unit sizes, y axis number of repeat units in the analysed dataset, whose size is given as Total bases. Peaks corresponding to the size of a 45S rDNA unit in particular chromosome arms are indicated by red arrows. In 5DS, the unit sizes need to be corrected by coefficient of 0.92 to adjust for concordance between the map and the sequence.
1BS
Total bases 305370 Mb
Repeat unit size (kb)
C
6BS
Total bases 117793 Mb
Repeat unit size (kb)
D
5DS
Total bases 480127 Mb
Repeat unit size (kb)

### Slide 3
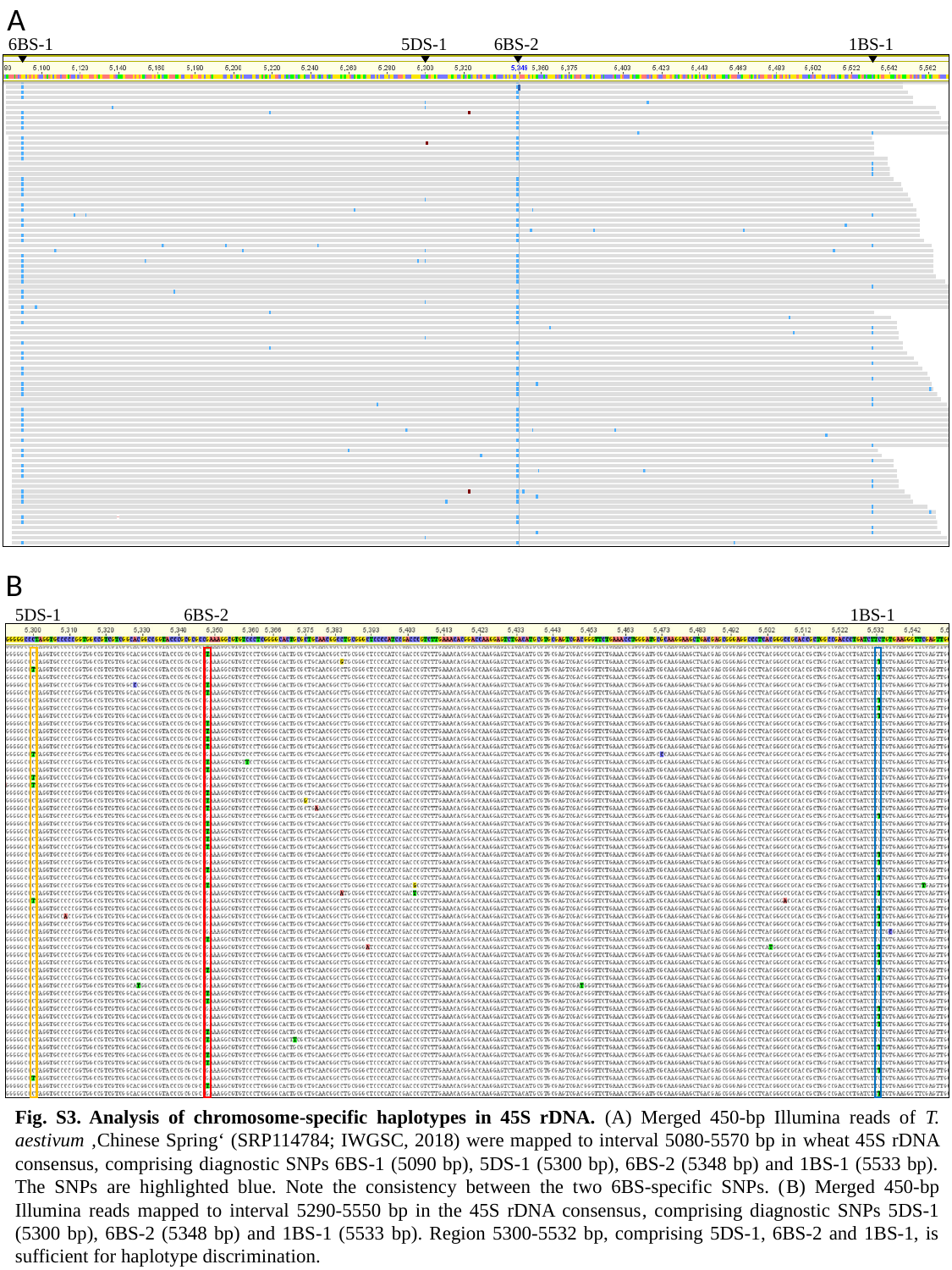

A
6BS-1
5DS-1
6BS-2
1BS-1
B
5DS-1
6BS-2
1BS-1
Fig. S3. Analysis of chromosome-specific haplotypes in 45S rDNA. (A) Merged 450-bp Illumina reads of T. aestivum ‚Chinese Spring‘ (SRP114784; IWGSC, 2018) were mapped to interval 5080-5570 bp in wheat 45S rDNA consensus, comprising diagnostic SNPs 6BS-1 (5090 bp), 5DS-1 (5300 bp), 6BS-2 (5348 bp) and 1BS-1 (5533 bp). The SNPs are highlighted blue. Note the consistency between the two 6BS-specific SNPs. (B) Merged 450-bp Illumina reads mapped to interval 5290-5550 bp in the 45S rDNA consensus, comprising diagnostic SNPs 5DS-1 (5300 bp), 6BS-2 (5348 bp) and 1BS-1 (5533 bp). Region 5300-5532 bp, comprising 5DS-1, 6BS-2 and 1BS-1, is sufficient for haplotype discrimination.

### Slide 4
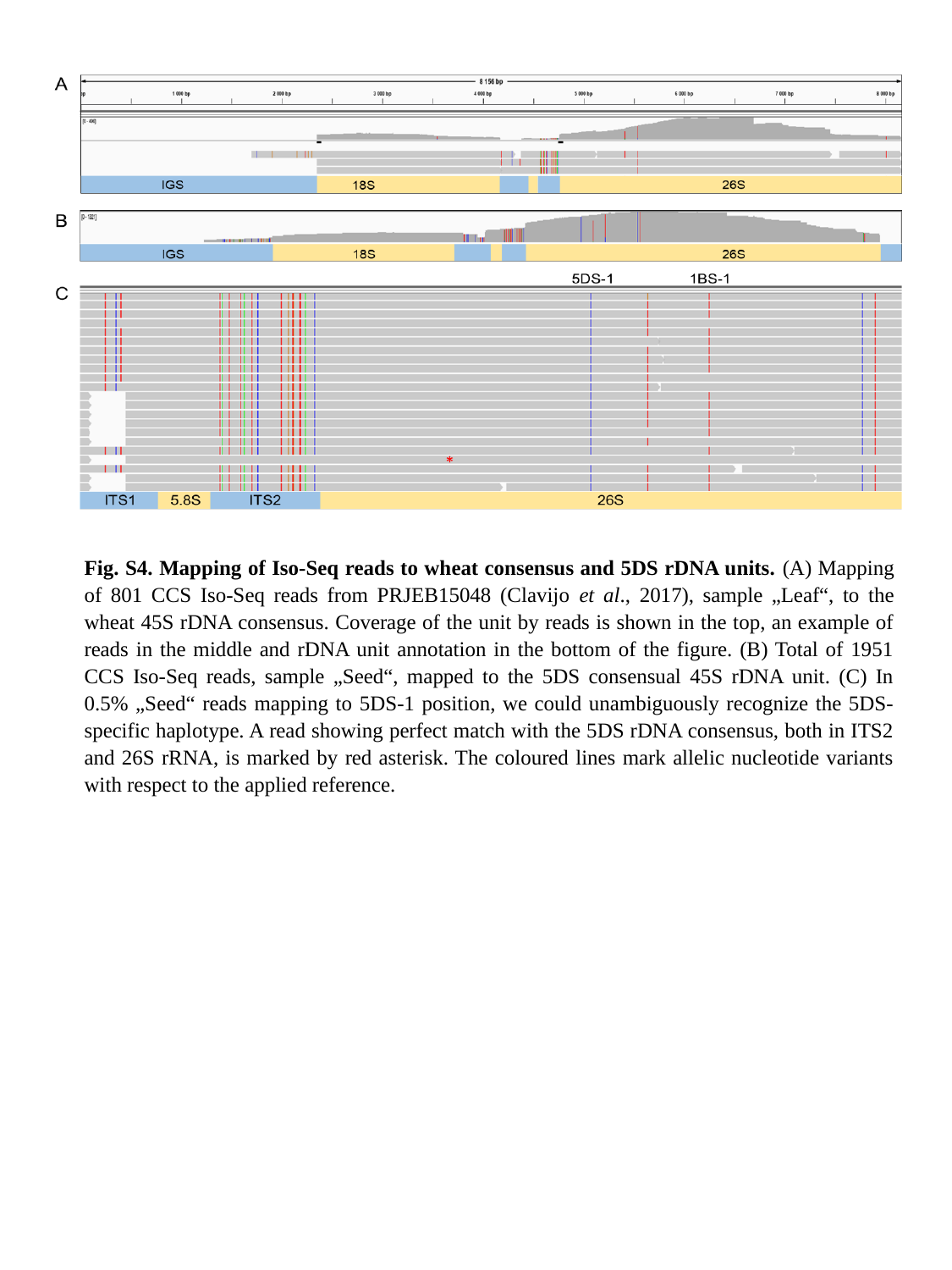

Fig. S4. Mapping of Iso-Seq reads to wheat consensus and 5DS rDNA units. (A) Mapping of 801 CCS Iso-Seq reads from PRJEB15048 (Clavijo et al., 2017), sample „Leaf“, to the wheat 45S rDNA consensus. Coverage of the unit by reads is shown in the top, an example of reads in the middle and rDNA unit annotation in the bottom of the figure. (B) Total of 1951 CCS Iso-Seq reads, sample „Seed“, mapped to the 5DS consensual 45S rDNA unit. (C) In 0.5% „Seed“ reads mapping to 5DS-1 position, we could unambiguously recognize the 5DS-specific haplotype. A read showing perfect match with the 5DS rDNA consensus, both in ITS2 and 26S rRNA, is marked by red asterisk. The coloured lines mark allelic nucleotide variants with respect to the applied reference.

### Slide 5
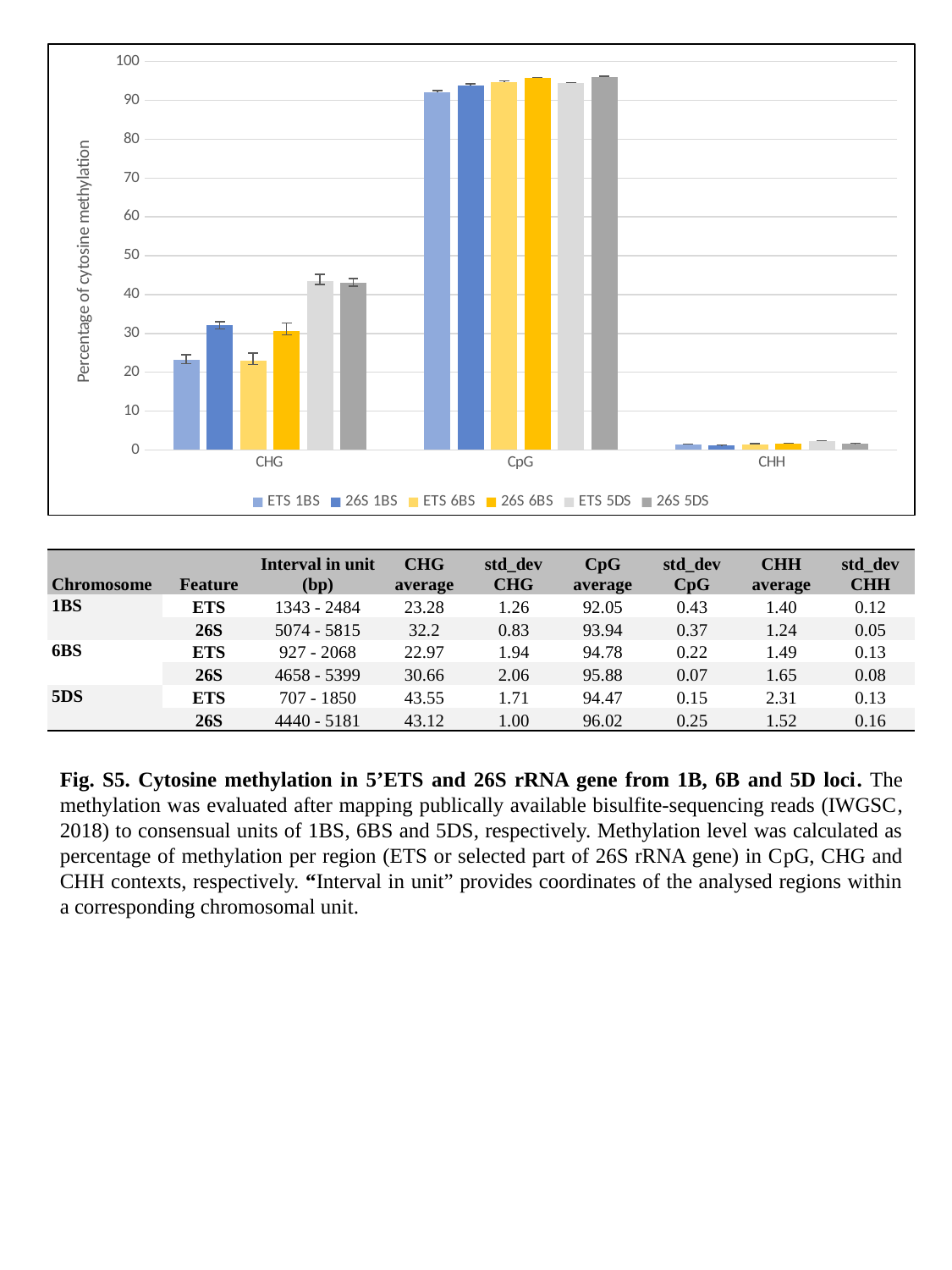

#### Chart
| Category | ETS | 26S | ETS | 26S | ETS | 26S |
|---|---|---|---|---|---|---|
| CHG | 23.281340916951468 | 32.2 | 22.969528198242134 | 30.66 | 43.550083160400334 | 43.12 |
| CpG | 92.05261993408199 | 93.94 | 94.77456156412757 | 95.88 | 94.46776580810543 | 96.02 |
| CHH | 1.3977168003718 | 1.24 | 1.4923009475072202 | 1.65 | 2.3139521280924438 | 1.52 || Chromosome | Feature | Interval in unit (bp) | CHG average | std\_dev CHG | CpG average | std\_dev CpG | CHH average | std\_dev CHH |
| --- | --- | --- | --- | --- | --- | --- | --- | --- |
| 1BS | ETS | 1343 - 2484 | 23.28 | 1.26 | 92.05 | 0.43 | 1.40 | 0.12 |
| | 26S | 5074 - 5815 | 32.2 | 0.83 | 93.94 | 0.37 | 1.24 | 0.05 |
| 6BS | ETS | 927 - 2068 | 22.97 | 1.94 | 94.78 | 0.22 | 1.49 | 0.13 |
| | 26S | 4658 - 5399 | 30.66 | 2.06 | 95.88 | 0.07 | 1.65 | 0.08 |
| 5DS | ETS | 707 - 1850 | 43.55 | 1.71 | 94.47 | 0.15 | 2.31 | 0.13 |
| | 26S | 4440 - 5181 | 43.12 | 1.00 | 96.02 | 0.25 | 1.52 | 0.16 |
Fig. S5. Cytosine methylation in 5’ETS and 26S rRNA gene from 1B, 6B and 5D loci. The methylation was evaluated after mapping publically available bisulfite-sequencing reads (IWGSC, 2018) to consensual units of 1BS, 6BS and 5DS, respectively. Methylation level was calculated as percentage of methylation per region (ETS or selected part of 26S rRNA gene) in CpG, CHG and CHH contexts, respectively. “Interval in unit” provides coordinates of the analysed regions within a corresponding chromosomal unit.
