## Supplemental Tables S1-S7 for "Anatomy, transcription dynamics and evolution of wheat ribosomal RNA loci deciphered by a multi-omics approach"

| **Table S1. Optical map statistics** | | |  |  |  |
| --- | --- | --- | --- | --- | --- |
|  | **Chromosome arm** | **1AS** | **1BS** | **6BS** | **5DS** |
| Input | **No. of arms sorted (million)** | 4 | 4 | 3.6 | 4.5 |
|  | **DNA amount (µg)** | 2.2 | 2.6 | 3 | 1.2 |
| Molecules | **Labelling chemistry** | NLRS | NLRS | NLRS | DLS |
|  | **Raw data ˃ 150 kb (Gb)** | 75 | 320 | 124 | 81 |
|  | **Filtered molecules N50 (kb)** | 209 | 197 | 207 | 227 |
|  | **Arm coverage** | 273x | 1019x | 298x | 315x |
| Assembly | **No. of contigs** | 300 | 369 | 566 | 35 |
|  | **Assembly length (Mb)** | 222 | 294 | 360 | 208 |
|  | **Contig N50 (Mb)** | 0.9 | 1.49 | 0.93 | 21 |
|  | **Average contig length (Mb)** | 0.74 | 0.8 | 0.64 | 5.85 |

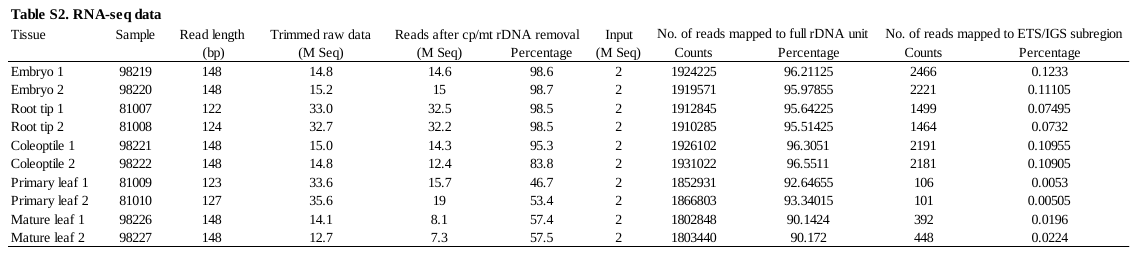

| Table S3. Frequency of allelic variants at diagnostic SNPs in Illumina reads of flow-sorted chromosome arms | | | | | | | | | | | | | |
| --- | --- | --- | --- | --- | --- | --- | --- | --- | --- | --- | --- | --- | --- |
|  | SNP | 6BS-1 (5090) | | 6BS-2 (5348) | | 1BS-1 (5533) | | 1BS-2 (8010) | | 5DS-1 (5300) | | 5DS-2 (8000) | |
|  | (position^1^) | No. | % | No. | % | No. | % | No. | % | No. | % | No. | % |
| **1AS** | A | 2 | 0.7 | 17 | 7.8 | 0 | 0.0 | 0 | 0.0 | 0 | 0.0 | 53 | 25.7 |
|  | C | 1 | 0.4 | 12 | 5.5 | 199 | 85.4 | 169 | 77.5 | 189 | 93.1 | 153 | 74.3 |
|  | T | 18 | 6.6 | 17 | 7.8 | 24 | 10.3 | 48 | 22.0 | 14 | 6.9 | 0 | 0.0 |
|  | G | 251 | 92.3 | 172 | 78.9 | 10 | 4.3 | 1 | 0.5 | 0 | 0.0 | 0 | 0.0 |
|  | - | 0 | 0.0 | 0 | 0.0 | 0 | 0.0 | 0 | 0.0 | 0 | 0.0 | 0 | 0.0 |
|  | Total | 272 |  | 218 |  | 233 |  | 218 |  | 203 |  | 206 |  |
| **1BS** | A | 12 | 0.1 | 31 | 0.3 | 13 | 0.1 | 11 | 0.1 | 3 | 0.0 | 17467 | 99.3 |
|  | C | 14 | 0.1 | 1 | 0.0 | 747 | 5.5 | 123 | 0.7 | 6424 | 99.8 | 106 | 0.6 |
|  | T | 69 | 0.4 | 45 | 0.4 | 12749 | 94.1 | 18256 | 99.0 | 3 | 0.0 | 6 | 0.0 |
|  | G | 16379 | 99.3 | 10075 | 99.2 | 25 | 0.2 | 41 | 0.2 | 3 | 0.0 | 9 | 0.1 |
|  | - | 17 | 0.1 | 4 | 0.0 | 10 | 0.1 | 6 | 0.0 | 5 | 0.1 | 8 | 0.0 |
|  | Total | 16491 |  | 10156 |  | 13544 |  | 18437 |  | 6438 |  | 17596 |  |
| **6BS** | A | 2 | 0.3 | 0 | 0.0 | 2 | 0.1 | 2 | 0.3 | 0 | 0.0 | 676 | 98.7 |
|  | C | 2 | 0.3 | 0 | 0.0 | 2982 | 99.5 | 738 | 97.7 | 55 | 100.0 | 6 | 0.9 |
|  | T | 714 | 94.3 | 346 | 98.6 | 11 | 0.4 | 14 | 1.9 | 0 | 0.0 | 2 | 0.3 |
|  | G | 39 | 5.2 | 5 | 1.4 | 1 | 0.0 | 1 | 0.1 | 0 | 0.0 | 0 | 0.0 |
|  | - | 0 | 0.0 | 0 | 0.0 | 1 | 0.0 | 0 | 0.0 | 0 | 0.0 | 1 | 0.1 |
|  | Total | 757 |  | 351 |  | 2997 |  | 755 |  | 55 |  | 685 |  |
| **5DS** | A | 2 | 0.2 | 3 | 0.5 | 0 | 0.0 | 0 | 0.0 | 0 | 0.0 | 74 | 13.3 |
|  | C | 0 | 0.0 | 0 | 0.0 | 763 | 95.9 | 540 | 97.1 | 47 | 13.9 | 484 | 86.7 |
|  | T | 110 | 9.0 | 45 | 7.7 | 32 | 4.0 | 16 | 2.9 | 290 | 86.1 | 0 | 0.0 |
|  | G | 1117 | 90.9 | 533 | 91.7 | 1 | 0.1 | 0 | 0.0 | 0 | 0.0 | 0 | 0.0 |
|  | - | 0 | 0.0 | 0 | 0.0 | 0 | 0.0 | 0 | 0.0 | 0 | 0.0 | 0 | 0.0 |
|  | Total | 1229 |  | 581 |  | 796 |  | 556 |  | 337 |  | 558 |  |
|  |  |  | 6BS variant | |  |  | 1BS variant | |  | 5DS variant | | Non-B variant | |
| ^1^ SNP position in the wheat 45S rDNA consensus | | | | | | |  |  |  |  |  |  |  |
| Chromosomal Illumina reads obtained from IWGSC, 2014 | | | | | | | |  |  |  |  |  |  |

**Table S4. Frequency of allelic variants at diagnostic SNPs in Illumina reads from three *T. aestivum* 'Chinese Spring' genome projects**

|  | SNP (position^1^) | 6BS-1 (5090) | | 6BS-2 (5348) | | 1BS-1 (5533) | | 1BS-2 (8010) | | 5DS-1 (5300) | | 5DS-2 (8000) | |
| --- | --- | --- | --- | --- | --- | --- | --- | --- | --- | --- | --- | --- | --- |
|  |  | No. | % | No. | % | No. | % | No. | % | No. | % | No. | % |
| **RefSeq v1.0^2^** | A | 16 | 0.0 | 74 | 0.2 | 19 | 0.1 | 31 | 0.1 | 13 | 0.0 | 33196 | 95.1 |
|  | C | 10 | 0.0 | 92 | 0.3 | 23647 | 64.5 | 21183 | 62.7 | 31952 | 95.7 | 1676 | 4.8 |
|  | T | 19583 | 55.9 | 18822 | 57.3 | 12951 | 35.3 | 12538 | 37.1 | 1385 | 4.2 | 10 | 0.0 |
|  | G | 15438 | 44.0 | 13886 | 42.2 | 48 | 0.1 | 27 | 0.1 | 23 | 0.1 | 23 | 0.1 |
|  | - | 0 | 0.0 | 1 | 0.0 | 0 | 0.0 | 1 | 0.0 | 0 | 0.0 | 0 | 0.0 |
|  | Total | 35047 |  | 32875 |  | 36665 |  | 33780 |  | 33373 |  | 34905 |  |
| **Triticum 3.1^3^** | A | 16 | 0.0 | 73 | 0.2 | 12 | 0.0 | 27 | 0.1 | 4 | 0.0 | 49039 | 95.3 |
|  | C | 12 | 0.0 | 74 | 0.2 | 31233 | 60.0 | 30399 | 58.9 | 44175 | 95.8 | 2354 | 4.6 |
|  | T | 28604 | 53.0 | 24443 | 53.9 | 20747 | 39.8 | 21135 | 41.0 | 1915 | 4.2 | 27 | 0.1 |
|  | G | 25301 | 46.9 | 20796 | 45.8 | 70 | 0.1 | 28 | 0.1 | 24 | 0.1 | 30 | 0.1 |
|  | - | 0 | 0.0 | 0 | 0.0 | 1 | 0.0 | 0 | 0.0 | 0 | 0.0 | 0 | 0.0 |
|  | Total | 53933 |  | 45386 |  | 52063 |  | 51589 |  | 46118 |  | 51450 |  |
| **TGACv1^4^** | A | 1 | 0.0 | 13 | 0.2 | 3 | 0.0 | 1 | 0.0 | 2 | 0.0 | 6263 | 94.9 |
|  | C | 1 | 0.0 | 0 | 0.0 | 5565 | 67.7 | 3894 | 59.5 | 6960 | 96.0 | 328 | 5.0 |
|  | T | 4083 | 56.2 | 4548 | 63.1 | 2638 | 32.1 | 2642 | 40.4 | 286 | 3.9 | 5 | 0.1 |
|  | G | 3180 | 43.8 | 2643 | 36.7 | 11 | 0.1 | 8 | 0.1 | 1 | 0.0 | 6 | 0.1 |
|  | - | 0 | 0.0 | 0 | 0.0 | 0 | 0.0 | 0 | 0.0 | 0 | 0.0 | 0 | 0.0 |
|  | Total | 7265 |  | 7204 |  | 8217 |  | 6545 |  | 7249 |  | 6602 |  |
|  |  |  | 6BS variant | |  |  | 1BS variant | |  | 5DS variant | | Non-B variant | |
| ^1^ | SNP position in the wheat 45S rDNA consensus | | | | |  |  |  |  |  |  |  |  |
| ^2^ | IWGSC, 2018 |  |  |  |  |  |  |  |  |  |  |  |  |
| ^3^ | Zimin *et al*., 2017 | |  |  |  |  |  |  |  |  |  |  |  |
| ^4^ | Clavijo *et al*., 2017 | |  |  |  |  |  |  |  |  |  |  |  |

**Table S5. Variant analysis of 26S rRNA in different tissues.** The four diagnostic SNPs were used to discriminate between transcription from 1BS and 6BS rDNA loci.

(A) **Frequency of diagnostic nucleotides in mapped rRNA-seq Illumina reads (in percentages)**

|  |  |  |  | **embryo** | | **coleoptile** | | **root tip** | | **primary leaf** | | **mature leaf** | |
| --- | --- | --- | --- | --- | --- | --- | --- | --- | --- | --- | --- | --- | --- |
| **Position** | **Type** | **Reference** | **Allele** | **98219** | **98220** | **98221** | **98222** | **81007** | **81008** | **81009** | **81010** | **98226** | **98227** |
| **5090** | 6BS-1 | G | T | 32.50 | 33.19 | 35.20 | 34.70 | 31.51 | 31.29 | 27.02 | 29.69 | 26.53 | 26.18 |
| **5090** |  | G | G | 66.81 | 66.20 | 64.13 | 64.70 | 68.14 | 68.45 | 72.59 | 69.97 | 72.82 | 73.18 |
| **5348** | 6BS-2 | G | T | 35.64 | 34.38 | 37.98 | 37.19 | 35.39 | 36.51 | 32.69 | 34.80 | 29.11 | 27.10 |
| **5348** |  | G | G | 63.69 | 64.68 | 61.12 | 62.27 | 63.51 | 62.58 | 66.14 | 64.59 | 70.02 | 72.01 |
| **5533** | 1BS-1 | C | T | 64.78 | 64.22 | 62.48 | 63.01 | 66.41 | 66.01 | 70.74 | 67.36 | 71.64 | 72.20 |
| **5533** |  | C | C | 34.33 | 34.77 | 36.61 | 36.00 | 33.33 | 33.67 | 28.96 | 32.36 | 27.36 | 26.73 |
| **8010** | 1BS-2 | C | T | 67.46 | 68.17 | 65.59 | 65.91 | 72.84 | 72.88 | 76.51 | 72.82 | 74.78 | 75.46 |
| **8010** |  | C | C | 32.54 | 31.83 | 34.41 | 34.09 | 27.15 | 27.11 | 23.49 | 27.18 | 25.25 | 24.54 |

^1^ SNP position in the wheat 45S rDNA consensus

The "T" allele was statistically evaluated by a Pearson chi square test (categories - "T" allele counts; total counts)

**(B) Statistical evaluation of SNP frequencies between the tissues**

| **Statistical evaluation of differences in variant representation at the 5090 position of the 26S rRNA gene** | | | | | | | | | | | | |
| --- | --- | --- | --- | --- | --- | --- | --- | --- | --- | --- | --- | --- |
|  |  | | **81007** | **81008** | **81009** | **81010** | **98219** | **98220** | **98221** | **98222** | **98226** | **98227** |
| Root tip | | **81007** | - | 0.456 | **171.461** | **27.923** | **9.086** | **26.58** | **121.058** | **90.851** | **255.332** | **303.8** |
| Root tip | | **81008** | 0.5141 | **-** | **149.552** | **20.632** | **48.906** | **32.638** | **129.99** | **99.387** | **223.376** | **267.137** |
| Primary leaf | | **81009** | **0.0001** | **0.0001** | - | 149.552 | **59.946** | **268.307** | **344.801** | **570.133** | **506.047** | 2.444 |
| Primary leaf | | **81010** | **0.0001** | **0.0001** | 0.0001 | - | **70.534** | **111.205** | **259.628** | **1004.195** | **33370.88** | **3621.379** |
| Embryo | | **98219** | **0.0026** | **0.0001** | **0.0001** | **0.0001** | - | 4.805 | **70.15** | **46.667** | **391.546** | **456.878** |
| Embryo | | **98220** | **0.0001** | **0.0001** | **0.0001** | **0.0001** | 0.0286 | - | **40.305** | **22.787** | **497.852** | **575.016** |
| Coleoptile | | **98221** | **0.0001** | **0.0001** | **0.0001** | **0.0001** | **0.0001** | **0.0001** | **-** | 2.373 | **793.182** | **894.841** |
| Coleoptile | | **98222** | **0.0001** | **0.0001** | **0.0001** | **0.0001** | **0.0001** | **0.0001** | 0.1234 | **-** | **708.192** | **802.6001** |
| Mature leaf | | **98226** | **0.0001** | **0.0001** | **0.0001** | **0.0001** | **0.0001** | **0.0001** | **0.0001** | **0.0001** | **-** | 1.553 |
| Mature leaf | | **98227** | **0.0001** | **0.0001** | **0.0001** | **0.0001** | **0.0001** | **0.0001** | **0.0001** | **0.0001** | 0.2128 | **-** |

| Footnote: |  |
| --- | --- |
| Above the diagonal - Pearson's chi square values | **In blue - moderately significant (P<0.05)** |
| Below the diagonal - p values | **In blue and underlined - highly significant (P<0.001)** |

| **Statistical evaluation of differences in variant representation at 5348 position within the 26S rRNA gene** | | | | | | | | | | | |
| --- | --- | --- | --- | --- | --- | --- | --- | --- | --- | --- | --- |
|  |  | **81007** | **81008** | **81009** | **81010** | **98219** | **98220** | **98221** | **98222** | **98226** | **98227** |
| Root tip | **81007** | **-** | 0.659 | 1.032 | 0.476 | 1.843 | 0.123 | **10.032** | **6.736** | **15.604** | **32.850** |
| Root tip | **81008** | 0.4170 | **-** | 1.032 | 0.476 | 1.843 | 0.247 | **5.320** | **2.972** | **22.957** | **43.241** |
| Primary leaf | **81009** | 0.3096 | 0.4170 | **-** | 3.356 | **6.453** | 2.068 | **20.149** | **15.142** | **15.142** | **9.853** |
| Primary leaf | **81010** | 0.4901 | 0.4901 | 0.067 | **-** | 3.356 | 0.121 | **7.219** | **4.248** | **24.328** | **46.584** |
| Embryo | **98219** | 0.1746 | 0.1746 | **0.011** | 0.4683 | **-** | 1.106 | 3.791 | 1.730 | **31.372** | **56.040** |
| Embryo | **98220** | 0.1746 | 0.6194 | 0.150 | 0.7275 | 0.2929 | **-** | **8.745** | **5.535** | **20.088** | **40.121** |
| Coleoptile | **98221** | **0.0015** | **0.0211** | **0.000** | **0.0072** | 0.0515 | **0.0031** | **-** | 0.4368 | **56.046** | **87.943** |
| Coleoptile | **98222** | **0.0095** | **0.0847** | **0.000** | **0.0393** | 0.1884 | **0.0186** | 0.5087 | **-** | **48.189** | **78.348** |
| Mature leaf | **98226** | **0.0001** | **0.0001** | **0.000** | **0.0001** | **0.0001** | **0.0001** | **0.0001** | **0.0001** | **-** | 3.330 |
| Mature leaf | **98227** | **0.0001** | **0.0001** | **0.002** | **0.0001** | **0.0001** | **0.0001** | **0.0001** | **0.0001** | 0.06800 | **-** |

| **Statistical evaluation of differences in variant representation at the 5533 position within the 26S rRNA gene** | | | | | | | | | | | |
| --- | --- | --- | --- | --- | --- | --- | --- | --- | --- | --- | --- |
|  |  | **81007** | **81008** | **81009** | **81010** | **98219** | **98220** | **98221** | **98222** | **98226** | **98227** |
| Root tip | **81007** | **-** | 0.4388 | 47.64 | 2.414 | 7.048 | 12.377 | **41.453** | **30.720** | **62.95** | **77.25** |
| Root tip | **81008** | 0.5077 | **-** | **58.679** | **5.083** | **4.091** | **8.402** | **34.256** | **24.479** | **75.129** | **90.889** |
| Primary leaf | **81009** | 0.0001 | 0.5077 | - | 30.579 | **92.091** | 107.173 | **179.570** | **155.826** | 1.862 | **4.877** |
| Primary leaf | **81010** | 0.1203 | **0.0241** | 0.0001 | - | **18.344** | **26.364** | **66.614** | **57.548** | **44.193** | **56.562** |
| Embryo | **98219** | 0.0079 | **0.0430** | **0.0002** | **0.0001** | **-** | 0.8040 | **14.408** | **8.428** | **110.509** | **129.344** |
| Embryo | **98220** | 0.0005 | 0.0037 | **0.0003** | **0.0001** | 0.370 | **-** | **8.103** | **3.874** | **126.088** | **145.887** |
| Coleoptile | **98221** | **0.0001** | **0.0001** | **0.0004** | **0.0001** | 0.0015 | **0.0044** | **-** | 0.782 | **200.250** | **225.567** |
| Coleoptile | **98222** | **0.0001** | **0.0001** | **0.0005** | **0.0001** | 0.0037 | 0.0490 | 0.3765 | **-** | **176.156** | **199.842** |
| Mature leaf | **98226** | **0.0001** | **0.0001** | 0.1724 | **0.0001** | **0.0001** | **0.0001** | **0.0001** | **0.0001** | **-** | 0.6363 |
| Mature leaf | **98227** | **0.0001** | **0.0001** | **0.0272** | **0.0001** | **0.0001** | **0.0001** | **0.0001** | **0.0001** | 0.4251 | **-** |

| **Statistical evaluation of differences in variant representation at the 8010 position within the 26S rRNA gene** | | | | | | | | | | | |
| --- | --- | --- | --- | --- | --- | --- | --- | --- | --- | --- | --- |
|  |  | **81007** | **81008** | **81009** | **81010** | **98219** | **98220** | **98221** | **98222** | **98226** | **98227** |
| Root tip | **81007** | **-** | 0.035 | **83.078** | 0.001 | **112.462** | **88.274** | **225.104** | **202.672** | **232,230** | **113.008** |
| Root tip | **81008** | 0.8525 | **-** | **75.811** | 0.023 | **108.731** | **85.981** | **224.876** | **193.754** | **64.751** | **101.413** |
| Primary leaf | **81009** | **0.0001** | **0.0001** | - | 85.920 | **425.466** | **372.387** | **617.018** | **581.7703** | **4.197** | 0.003 |
| Primary leaf | **81010** | 0.9692 | 0.8788 | 0.0001 | - | **22.860** | **120.837** | **94.670** | **241.430** | **76.631** | **6.503** |
| Embryo | **98219** | **0.0001** | **0.0001** | **0.0001** | **0.0001** | **-** | 1.538 | **31.041** | **20.957** | **542.940** | **653.752** |
| Embryo | **98220** | **0.0001** | **0.0001** | **0.0001** | **0.0001** | 0.2147 | **-** | **43.925** | **32.148** | **459.757** | **561.219** |
| Coleoptile | **98221** | **0.0001** | **0.0001** | **0.0001** | **0.0001** | **0.0001** | **0.0001** | **-** | 1.0212 | **891.659** | **951.220** |
| Coleoptile | **98222** | **0.0001** | **0.0001** | **0.0001** | **0.0001** | **0.0001** | **0.0001** | 0.3121 | **-** | **771.025** | **897.922** |
| Mature leaf | **98226** | **0.0001** | **0.0001** | **0.0405** | **0.0001** | **0.0001** | **0.0001** | **0.0001** | **0.0001** | **-** | 1.8186 |
| Mature leaf | **98227** | **0.0001** | **0.0001** | 0.9559 | **0.0108** | **0.0001** | **0.0001** | **0.0001** | **0.0001** | 0.1775 | **-** |

| Table S6. Transcripts per kilobase million (TPM) values for nuclear, chloroplast and mitochondrial 16S/18S and 23/26S rRNAs across five tissues | | | | | | | | | | |
| --- | --- | --- | --- | --- | --- | --- | --- | --- | --- | --- |
|  | embryo | | coleoptile | | root tip | | primary leaf | | mature leaf | |
|  | 98219 | 98220 | 98221 | 98222 | 81007 | 81008 | 81009 | 81010 | 98226 | 98227 |
| nuclear 18S | 494502 | 483613 | 471110 | 484340 | 546540 | 550230 | 275920 | 299541 | 271150 | 264829 |
| nuclear 26S | 446378 | 449965 | 434594 | 438905 | 412718 | 408399 | 202510 | 231254 | 233223 | 234483 |
| chloroplast 16S | 788 | 1120 | 12005 | 7635 | 683 | 624 | 133758 | 123490 | 132817 | 131103 |
| chloroplast 23S | 905 | 1414 | 14145 | 8800 | 540 | 493 | 141222 | 126463 | 151508 | 156115 |
| mitochondrial 18S | 2973 | 2626 | 2377 | 2262 | 1283 | 1478 | 535 | 571 | 1202 | 1144 |
| mitochondrial 26S | 2922 | 2608 | 2100 | 2069 | 1617 | 1889 | 655 | 638 | 1053 | 1016 |

| **Table S7. Cytosine methylation in 5’ETS of 45S rDNA units - statistics** | | | | | | |
| --- | --- | --- | --- | --- | --- | --- |
|  | **CHG_replica1** | **CHG_replica2** | | **CHG_replica3** | | **Average** |
| rDNA_1BS | 22.04 | 23.25 | | 24.55 | | 23.28 |
| rDNA_5DS | 41.66 | 44.01 | | 44.98 | | 43.55 |
| rDNA_6BS | 20.94 | 23.16 | | 24.80 | | 22.97 |
|  | **CHH_replica1** | **CHH_replica2** | | **CHH_replica3** | | **Average** |
| rDNA_1BS | 1.26 | 1.47 | | 1.46 | | 1.40 |
| rDNA_5DS | 2.19 | 2.44 | | 2.31 | | 2.31 |
| rDNA_6BS | 1.34 | 1.59 | | 1.55 | | 1.49 |
|  | **CpG_replica1** | **CpG_replica2** | | **CpG_replica3** | | **Average** |
| rDNA_1BS | 91.64 | 92.02 | | 92.50 | | 92.05 |
| rDNA_5DS | 94.50 | 94.30 | | 94.60 | | 94.47 |
| rDNA_6BS | 94.53 | 94.86 | | 94.94 | | 94.77 |
| **Independent two-tailed T-test comparing two chromosomal units at the time** | | | | | | |
| **CHG** |  |  |  | |  | |
| 1BS - 5DS | The t-value is -16.56881. The p-value is .000039. The result is significant at p < .05. | | | | | |
| 1BS - 6BS | The t-value is 0.23312. The p-value is .827113. The result is not significant at p < .05 | | | | | |
| 5DS - 6BS | The t-value is 13.81285. The p-value is .000159. The result is significant at p < .05. | | | | | |
| **CHH** |  |  |  | |  | |
| 1BS - 5DS | The t-value is -9.25979. The p-value is .000756. The result is significant at p < .05 | | | | | |
| 1BS - 6BS | The t-value is -0.9004. The p-value is .418815. The result is not significant at p < .05 | | | | | |
| 5DS - 6BS | The t-value is 7.73608. The p-value is .001504. The result is significant at p < .05. | | | | | |
| **CpG** |  |  |  | |  | |
| 1BS - 5DS | The t-value is -9.11654. The p-value is .000803. The result is significant at p < .05. | | | | | |
| 1BS - 6BS | The t-value is -9.72635. The p-value is .000626. The result is significant at p < .05. | | | | | |
| 5DS - 6BS | The t-value is -1.99513. The p-value is .116764. The result is not significant at p < .05. | | | | | |
